# Long-term effects of network-based fMRI neurofeedback training for sustained attention

**DOI:** 10.1101/2021.10.27.465722

**Authors:** Gustavo Santo Pedro Pamplona, Jennifer Heldner, Robert Langner, Yury Koush, Lars Michels, Silvio Ionta, Carlos Ernesto Garrido Salmon, Frank Scharnowski

**Affiliations:** Sensory-Motor Laboratory (SeMoLa), Jules-Gonin Eye Hospital/Fondation Asile des Aveugles, Department of Ophthalmology/University of Lausanne, Lausanne, Switzerland; InBrain Lab, Department of Physics, University of Sao Paulo, Ribeirao Preto, Brazil; Department of Psychiatry, Psychotherapy and Psychosomatics, Psychiatric Hospital, University of Zurich, Switzerland; Rehabilitation Engineering Laboratory (RELab), Department of Health Sciences and Technology, ETH Zurich, Zurich, Switzerland; Institute of Systems Neuroscience, Heinrich Heine University Dusseldorf, Dusseldorf, Germany; Institute of Neuroscience and Medicine, Brain & Behaviour (INM -7), Research Centre Julich, Julich, Germany; Department of Radiology and Biomedical Imaging, Yale School of Medicine, Yale University, New Haven, CT, USA; Department of Neuroradiology, University Hospital Zurich, Zurich, Switzerland; Neuroscience Center Zurich, University of Zurich and Swiss Federal Institute of Technology, Zurich, Switzerland; Zurich Center for Integrative Human Physiology (ZIHP), University of Zurich, Zurich, Switzerland; Department of Cognition, Emotion, and Methods in Psychology, Faculty of Psychology, University of Vienna, Vienna, Austria

**Author notes:** Both authors contributed equally (Carlos Ernesto Garrido Salmon, Frank Scharnowski).

**Keywords:** fMRI-neurofeedback, follow-up, sustained attention, functional connectivity, resting state, psychophysiological interaction (PPI), behavioral effects

## Abstract

Neurofeedback allows for learning voluntary control over one’s own brain activity, aiming to enhance cognition and clinical symptoms. A recent study improved sustained attention temporarily by training healthy participants to up-regulate the differential activity of the sustained attention network (SAN) minus the default mode network (DMN). However, long-term learning effects of functional magnetic resonance imaging (fMRI) neurofeedback training remain under-explored. Here, we evaluate the effects of network-based fMRI neurofeedback training for sustained attention by assessing behavioral and brain measures before, one day after, and two months after training. The behavioral measures include task as well as questionnaire scores, and the brain measures include activity and connectivity during self-regulation runs without feedback (i.e., transfer runs) and during resting-state runs. Neurally, we found that participants maintained their ability to control the differential activity during follow-up sessions. Further, exploratory analyses showed that the training-induced increase in FC between the DMN and occipital gyrus was maintained during follow-up transfer runs, but not during follow-up resting-state runs. Behaviorally, we found that enhanced sustained attention right after training returned to baseline level during follow-up. The discrepancy between lasting regulation-related brain changes but transient behavioral and resting-state effects raises the question of how neural changes induced by neurofeedback training translate to potential behavioral improvements. Since neurofeedback directly targets brain measures to indirectly improve behavior long-term, a better understanding of the brain-behavior associations during and after neurofeedback training is needed to develop its full potential as a promising scientific and clinical tool.

**Key points:**

- Participants were still able to self-regulate the differential activity between large-scale networks two months after the end of neurofeedback training and this during transfer runs without feedback.
- Lasting brain changes were also observed in the functional connectivity of trained regions in runs during which participants engaged in active self-regulation as well as during resting-state runs without concomitant self-regulation.
- The increased sustained attention we observed right after the end of neurofeedback training did not persist two months later.

## 1. INTRODUCTION

Neurofeedback is a form of biofeedback that provides individuals with real-time sensory information from their own brain activity, over which voluntary control can be learned with training [Sitaram et al., 2017]. Neurofeedback training has been associated with behavioral changes, which makes it an interesting approach for studying brain-behavior relationships [Sitaram et al., 2017; Sulzer et al., 2013a]. Neurofeedback training has also produced clinical benefits, which makes it a promising clinical intervention for the treatment of neurological and psychiatric disorders (e.g. Linhartová et al., 2019; Martz et al., 2020; Sokunbi, 2017; Sulzer et al., 2013; Taschereau-Dumouchel et al., 2022; Wang et al., 2018). The reported effects of neurofeedback training include transient as well as lasting changes. For cognitive enhancement and clinical applications of neurofeedback training, lasting effects are particularly important. Behaviorally, several studies reported that neurofeedback was associated with changes that lasted beyond the initial training [Amano et al., 2016; Mehler et al., 2018; Zilverstand et al., 2015a]. Some studies even found that clinical symptoms “continue to improve for weeks after neurofeedback” training [Rance et al., 2018]. Also neurally, lasting plastic brain changes have been reported, including resting-state functional connectivity (FC) [Megumi et al., 2015; Scheinost et al., 2013; Yuan et al., 2014; Zhang et al., 2013] and brain structural changes [Marins et al., 2019; Sampaio-Baptista et al., 2021]. Lasting brain changes combined with behavioral modulations induced by neurofeedback training provide new insights into how they relate to each other. Hence, investigating long-term effects of neurofeedback training will help understanding its learning mechanisms and might facilitate the use of neurofeedback for enhancing cognition and clinical symptoms.

Here we investigate lasting behavioral and neural changes following neurofeedback training of sustained attention. Sustained attention is a cognitive function that supports the continuous focus on a particular external object for extended periods of time. Neuroimaging correlates of sustained attention comprise the sustained attention network (SAN) [Langner and Eickhoff, 2013], which combines regions from the frontoparietal control network [Dosenbach et al., 2008] and the dorsal attention network (DAN) [Yeo et al., 2011]. In contrast, default mode network (DMN) activation is related to internally-focused cognitive processes and mind-wandering [Andrews-Hanna et al., 2014; Raichle et al., 2001]. DMN activation is therefore associated with stimulus-independent thoughts and reduced attention during the execution of an externally-oriented task [Hinds et al., 2011; Lawrence et al., 2003; Thompson et al., 2013; Weissman et al., 2006]. The SAN (more specifically, its DAN components [Spreng, 2012]) and DMN are intrinsically anticorrelated, as they are engaged in antagonistic processes reflecting externally- vs. internally-oriented attention [Fox et al., 2005; Spreng, 2012]. We recently demonstrated that sustained attention can be improved to some extent through training simultaneous up-regulation of the sustained attention network (SAN) and down-regulation of the default-mode network (DMN) using fMRI neurofeedback [Pamplona et al., 2020a]. We found that participants in the neurofeedback group were able to regulate their differential SAN-DMN activity and showed improved sustained attention directly after the training. No such improvement was observed in a test-retest control group, which only performed the behavioral sustained attention tasks, but did not undergo neurofeedback training.

Regarding lasting effects, we hypothesize that the neural and behavioral changes induced by the neurofeedback training persist beyond the training. Specifically, we hypothesized that regulation performance in brain regions successfully trained using neurofeedback and the associated improved sustained attention would be maintained long-term. We also hypothesized that functional connectivity changes specific to the successfully trained brain regions would be observed in the long term. To test these hypotheses, we analyzed unpublished data of runs without feedback (i.e., transfer runs) and resting-state runs, from before, immediately after, and two-months after neurofeedback training. We also explored the data in terms of immediate and lasting whole-brain activation and functional connectivity changes. Finally, we explored associations between brain connectivity changes and behavioral effects. Specifically, we investigated (i) the persistence of learned regulation in SAN, DMN, and their constituent regions two months after training to characterize maintained self-regulation; (ii) changes in pre-training functional connectivity directly after training and two months later to investigate lasting brain connectivity alterations with (transfer runs) and without regulation (resting-state runs); (iii) changes in resting-state functional connectivity of SAN and DMN regions directly after training and two months later using a graph theoretical approach; (iv) persistence of training-induced attention measured by task and questionnaires two months after neurofeedback training to assess the permanence of behavioral effects arising from neurofeedback training; and (v) associations of functional connectivity changes in transfer and resting-state runs with behavioral changes directly after training and two months later.

## 2. MATERIALS AND METHODS

### 2.1. Participants

We included data from a previously published study [Pamplona et al., 2020a] that comprised a neurofeedback training group who performed sustained attention tasks before and after training. The study also included a control group who only performed the sustained attention tasks twice without neurofeedback training, separated by a two-week interval which corresponds to the duration of the neurofeedback training in the experimental group. For the present study, we analyzed data only from the neurofeedback group, which consisted of 15 healthy volunteers (5 females, mean age: 27.9 ± 3.3 years old, age range = [22.6, 34.5] years old). Data included psychometric tasks, transfer and resting-state runs before, directly after, and two months after neurofeedback training. Exclusion criteria were left-handedness, strong vision deficiency that could not be corrected using contact lenses, insufficient knowledge of English, history of mental and/or cardiovascular disorders, not being able to abstain from alcohol or other drugs during the days of the experiment, and MRI contraindications. This study was approved by the local ethics committee of the Canton of Zurich in Switzerland. All participants read and signed the informed consent in accordance with the Declaration of Helsinki (2013) before taking part in the study. They received financial compensation of 25 CHF per hour for their participation.

### 2.2. Experimental procedure

#### Timeline of experimental procedure

Each participant took part in a five-day longitudinal study (Fig. 1) that involved fMRI-neurofeedback training and pre/post-training sessions for neural and behavioral assessment of neurofeedback training. The neurofeedback training consisted of ten real-time fMRI runs that took place on the second and third days of the experiment (Fig. 1), split into five runs each day. Each neurofeedback training session lasted about 45 min. The interval between training days for each participant was a maximum of seven days. Neurofeedback training runs consisted of five cycles of baseline, regulation, and intermittent feedback blocks, lasting 30 s, 40 s, and 4 s, respectively. To indicate the period of baseline, regulation, and feedback blocks, participants were presented with a black square, a black up-arrow, and a graded blue-to-red thermometer on the center of a white screen, respectively. We acquired individual transfer runs – used to test learned self-regulation in situations where feedback is not available – directly before and directly after training (second and third days of the experiment, respectively), as well as two transfer runs two months after the end of training (fifth day of the experiment). On the first, fourth, and fifth days of the experiment, the participants filled attention-related questionnaires (see section 2.3) and underwent anatomical and resting-state fMRI acquisitions. Attention tasks were performed outside the scanner. These measurements were made approximately at the same time of the day. The intervals between first and second days, between third and fourth days, and third and fifth days were 7 days maximum, 1 day maximum, and 61 ± 3 days, respectively. Hence, the terms pre-training, post-training, and follow-up correspond to measurements acquired on the first and second days, third and fourth days, and fifth days of the experiment, respectively.

**Figure 1.**
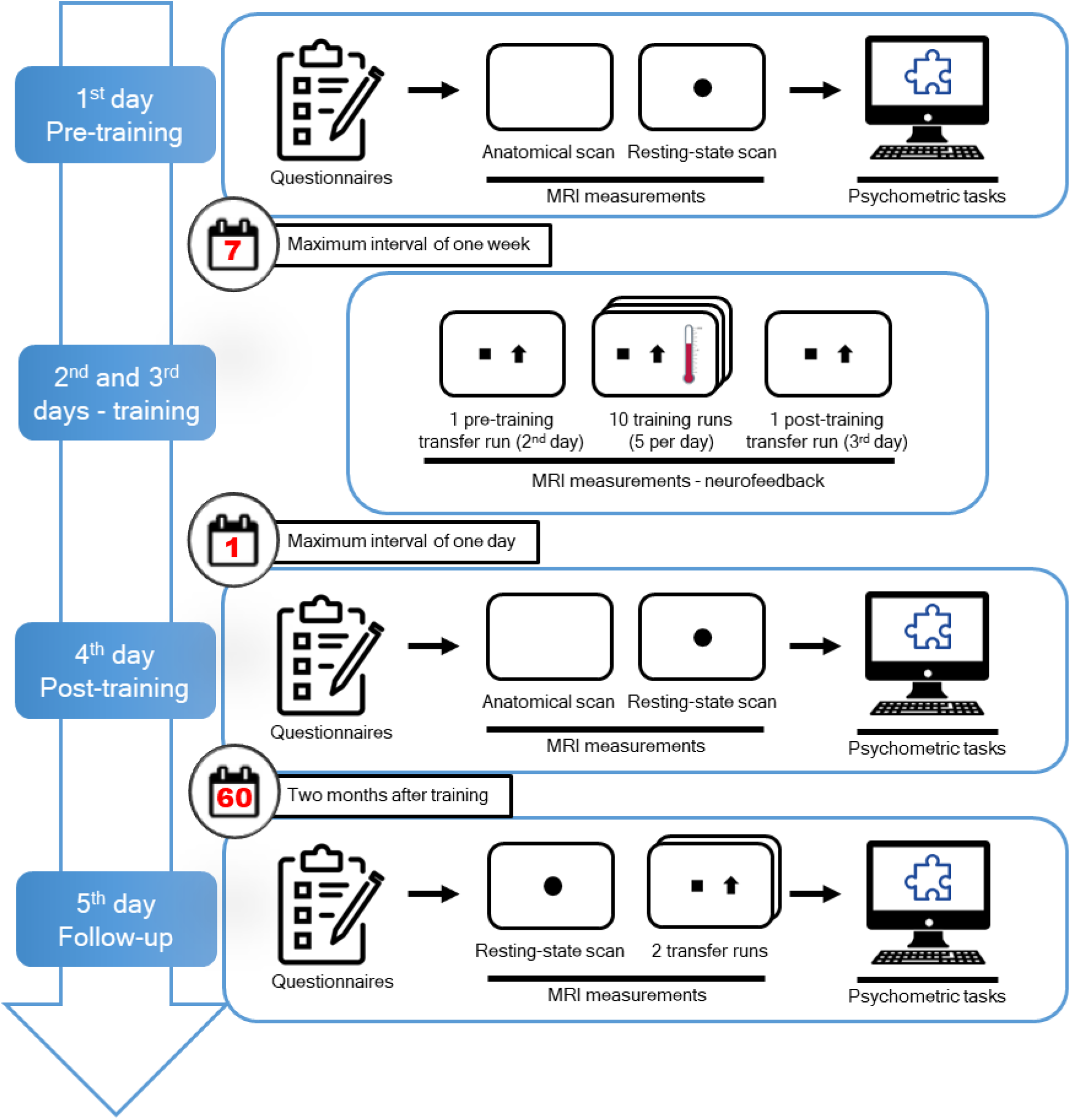
Timeline of the five-day neurofeedback experimental procedure. On the first, fourth, and fifth days of the experiment, participants underwent resting-state fMRI acquisitions and completed attention-related questionnaires (DSSQ and CFQ) as well as psychometric tasks on a computer (Continuous Performance Task, Switcher; PVT, Mental Rotation Task and Attentional Network Test). The neurofeedback training sessions occurred on the second and third days of the experiment and the participants performed self-regulation without feedback during transfer runs directly before and directly after training. Additionally, participants also performed self-regulation without feedback on two transfer runs on the fifth day. The five visits were conducted with a maximum of one week between the first and second days, one day between the third and fourth days, and two months between the third and fifth days.

#### Instructions

Instructions for self-regulation strategies during neurofeedback training were provided in written form outside the MR scanner room, prior to scanning. We instructed participants to relax and let their minds wander during baseline blocks and to engage in one of the suggested regulation strategies ((1) constantly reorienting the focus on different aspects of the arrow every 5-10 s; (2) focusing on the black up-arrow and bringing attention back to it whenever detecting task-unrelated thoughts; (3) staying in a state of high alertness) during regulation blocks. Participants were told that they could explore other regulation strategies and adopt the ones that worked best for them. Participants were also explicitly informed that, during baseline blocks, they should not plan regulation tasks. For the pre-training transfer run, participants were asked to choose one of the suggested regulation strategies and employ it during this run. For the post-training and follow-up transfer runs, participants were asked to use the strategy that worked best throughout the neurofeedback training.

#### MRI acquisition

All MR images were acquired on a Philips Achieva 3T MRI scanner with a 32-channel head coil in the MR center of the Psychiatric Hospital, University of Zürich, Switzerland. Functional images were acquired using a T2*-weighted gradient-echo-planar (EPI) sequence with repetition time/echo time (TR/TE) = 2000/30 ms, flip angle = 80°, and field of view (FOV) = 240 × 240 mm^2^. 37 slices were acquired in ascending order to cover the entire cerebrum (voxel size = 3 × 3 × 4 mm^3^, gap = 0.5 mm). SofTone mode was activated to reduce acoustic scanner noise. Anatomical T1-weighted brain images were acquired using a 3D MPRAGE (magnetization prepared gradient echo) sequence, TR/TE = 7.2/3.4 ms, 170 slices, voxel size = 1 × 1 × 1 mm^3^, flip angle = 8°, FOV = 240 mm x 240 mm^2^, duration = 3.5 min. Resting-state fMRI acquisitions comprised 200 scans (6 min 40 s) during which participants were asked to not move, relax, breath regularly, look at a central black circle presented on a white screen for visual fixation, and not to think about anything in particular. Neurofeedback training and transfer acquisitions comprised 190 scans (6 min 20 s) and 180 scans (6 min), respectively. Before every functional acquisition, five dummy scans were acquired to establish steady-state magnetizations. Visual stimuli were presented with MR-compatible goggles (Resonance Technology Inc.).

#### Definitions of target networks

To improve sustained attention through neurofeedback training, we simultaneously promoted the activation of four representative regions-of-interest (ROIs) from the sustained attention network (SAN) and the deactivation of four representative ROIs from the default mode network (DMN), areas positively and negatively associated with sustained attention performance, respectively. The SAN ROIs were defined using a mask of meta-analytic clusters from a comprehensive study on sustained attention [Langner and Eickhoff, 2013] (Table S1). The selected SAN ROIs were the anterior mid-cingulate cortex (aMCC), the right inferior frontal junction (R IFJ), the right temporoparietal junction (R TPJ), and the right intraparietal sulcus (R IPS), chosen to represent multiple functional aspects of the ability of sustained attention. The aMCC is related to conflict processing, monitoring performance, and enhanced vigilance [Hinds et al., 2013; Langner and Eickhoff, 2013; Weissman et al., 2006]; the R IFJ is related to stimuli discrimination and attention switching [Langner and Eickhoff, 2013]; the R TPJ is associated with bottom-up attention reorienting [Corbetta and Shulman, 2002; Weissman et al., 2006]; and the R IPS is associated with top-down attention reorienting [Corbetta and Shulman, 2002; Harris et al., 2000]. The ROIs representing the aMCC and the R IFJ were spatially eroded from the original meta-analytic clusters to reduce their volume. The selected DMN ROIs were the posterior cingulate cortex (PCC), medial prefrontal cortex (mPFC), and bilateral angular gyri (L Ang and R Ang). These regions are the most consistently reported DMN regions – the so-called core regions of the DMN – and robustly activated during self-generated tasks [Andrews-Hanna et al., 2014], in contrast to externally-oriented attention tasks. To account for individual differences, the DMN ROIs were defined using the resting-state acquisitions from each participant. More specifically, we first performed an independent component analysis (ICA) as implemented in Gift (mialab.mrn.org/software/gift) with a predefined number of 30 components. Next, using the Personode toolbox (Pamplona et al., 2020b; www.nitrc.org/projects/personode), we created 6-mm-radius spherical ROIs centered on probabilistic peaks that maximally represented each DMN regions for each individual (Table S1).

#### Feedback estimation and presentation

Real-time signal processing was performed using OpenNFT [Koush et al., 2017]. Before each real-time fMRI session, the MNI (Montreal Neurological Institute)-based ROIs were transformed into the current native space using SPM12 (www.fil.ion.ucl.ac.uk). First, for neurofeedback training runs, the signal averaged within each ROI was rescaled in real-time [Koush et al., 2012; Pamplona et al., 2020a; Scharnowski et al., 2012]. Next, the resulting signals were averaged within SAN and DMN separately. Finally, the difference between SAN and DMN signals (differential activity SAN minus DMN) was fed back intermittently to the participant as the thermometer level, right after regulation blocks. Participants were asked to raise the thermometer level as much as possible, which could be achieved either by SAN upregulation, DMN downregulation, or both. The thermometer level, comprised of 10 negative (for DMN > SAN), zero, and 10 positive readings (for SAN > DMN), was proportional to the participant’s performance in the current block. Feedback presentation was adaptive for each run based on performance in previous runs, i.e., feedback was made more difficult if the task was relatively easy for the participant and vice-versa. At the end of each run, a monetary reward proportional to their performance in each run was shown to the participant (CHF 20.6 ± 5.4 in total per participant) and added to the final compensation to the participation.

### 2.3. Psychometric tasks and questionnaires

To evaluate mental strategies associated with neurofeedback training, we asked participants to report the used strategies immediately after each neurofeedback training run. In addition, at the end of training and transfer runs, participants rated their level of concentration on the previous run on a scale ranging from 1 (very low) to 10 (very high). Self-reported concentration ratings from two participants were not collected due to technical issues with the communication system.

At the beginning of the first, fourth, and fifth days, participants also completed attention questionnaires [Cognitive Failure Questionnaire (CFQ) [Broadbent et al., 1982]] and their current state of attentiveness and stress in real-life situations [Dundee State Questionnaire (DSSQ) [Helton, 2004]]. Technical failures in the acquisition led to incomplete data collection: inclusion of 14-15 participants in the pre-training session, 6-7 participants in the post-training session, and 12-15 participants in the follow-up session (the number of participants varies depending on missing data specific to the sub-score).

At the end of the first, fourth, and fifth days, participants performed five attention-related tasks, as implemented in the Psychology Experiment Building Language (PEBL) software [Mueller and Piper, 2014], outside the scanner (Fig. 1). Attention tests were performed on a dedicated computer and in a separate experimental room with constant luminosity and noise (participants were asked to use earplugs). The selected tasks from PEBL were: (1) Continuous Performance Task (CPT) [Conners et al., 2003; Ogg et al., 2008; Piper et al., 2016], a go/no-go task designed to measure the sustained ability to either execute or withhold a speeded response; (2) Task-Switching Performance (Switcher) [Anderson, et al., 2012], designed to evaluate the cognitive flexibility in reorienting attention to switching rules; (3) Psychomotor Vigilance Test (PVT) [Dinges and Powell, 1985; Helton et al., 2007; Loh et al., 2004], designed to measure the level of alertness and its maintenance over time (sustained attention); (4) Mental Rotation Task [Berteau-Pavy et al., 2011; Shepard and Metzler, 1971], designed to evaluate the visual imagery ability in transforming spatial characteristics of an image; (5) Attentional Network Test (ANT) [Fan et al., 2002], designed to provide measurements of different facets of attention: phasic alerting, endogenous spatial orientating, and conflict resolution. The tasks were presented always in the same order. To avoid fatigue, there were 5-min breaks between the 2nd and 3rd tests and between the 4th and 5th tests.

### 2.4. Data analysis

Functional images from the transfer and resting-state runs as well the anatomical images were preprocessed using SPM12 in MATLAB (The MathWorks, Natick, MA, USA). First, functional images were slice-time corrected using the middle slice as reference. Then, three translation and three rotation parameters of head motion were estimated, and the functional images were spatially realigned to a created mean functional image. Next, the anatomical image was coregistered to the mean functional image and then segmented into tissue probability masks for gray matter, white matter, and cerebrospinal fluid (CSF) compartments. During the segmentation process, a deformation field was created, which was used to normalize the anatomical and functional images to the standard MNI template. Finally, the normalized functional images from the transfer runs were spatially smoothed using a Gaussian kernel of 8 mm full-width at half maximum (FWHM), and the normalized functional images from the resting-state runs were smoothed with a kernel of 6-mm FWHM.

#### 2.4.1. Transfer run activity and regulation-specific functional connectivity analyses

##### First-level analysis of transfer runs

We investigated differences in training-induced neural activity changes across pre-training, post-training, and follow-up transfer runs. For the first-level analysis, we specified for each run a general linear model (GLM) with two regressors of interest representing regulation and baseline conditions, and six covariates representing head motion. Regressors of interest were modeled as boxcar functions and convolved with the canonical hemodynamic response function implemented in SPM12. Next, beta values (regression weights) of regulation and baseline blocks for each participant and run were estimated voxel-wise. Contrasts were created for the activation differences between regulation and baseline blocks for each participant and run.

##### Long-term effects of regulation in trained networks

To examine the follow-up effects in brain self-regulation after neurofeedback training, we investigated whether the differential SAN-DMN activity, as well as activations within the SAN and the DMN and their constituent ROIs, differed between follow-up and the pre-training transfer runs. First, contrast values (regulation vs. baseline) were extracted using MarsBaR (marsbar.sourceforge.net, Brett, Anton, Valabregue, & Poline, 2002). Then, we averaged the contrasts from the four SAN and the four DMN ROIs to compute the SAN and DMN contrasts for each session, respectively, as well as the differential SAN-DMN signal. The contrasts from the two follow-up transfer runs were collapsed together. We then compared the differential SAN-DMN signal, as well as the contrasts for SAN and DMN and for their constituent ROIs separately, across follow-up and pre-training sessions using paired t-tests using RStudio (www.rstudio.com). The normality of each run-specific distribution was verified using Shapiro-Wilk tests. Statistical tests of the comparison of activity during follow-up compared to pre-training sessions were one-tailed because we hypothesized more positive estimates for differential SAN-DMN activity difference and for SAN activity, as well as more negative estimates for the DMN activity. We also estimated the effect sizes of the follow-up minus pre-training differences using Cohen’s d.

##### Long-term effects of regulation across the whole brain

First, individual contrast maps (regulation vs. baseline) for each session (i.e., pre-training, post-training, and follow-up) were entered into a second-level analysis in which subjects were treated as random effects. Then, voxel-wise one-sample t-tests were performed to map the group activations and deactivations for each session. We also created statistical maps comparing post- vs. pre-training sessions and follow-up vs. pre-training sessions. These statistical maps were obtained by entering individual contrast maps (post-minus pre-training or follow-up minus pre-training) as random effects in one-sample t-tests (which is equivalent to paired t-tests with partitioned errors [Henson, 2015]). All resulting group-level maps were submitted to the threshold-free cluster estimation (TFCE) approach (voxel-level threshold of p < 0.001 uncorrected for multiple comparisons, 10000 permutations). This approach provides high sensitivity for detecting both large and small clusters [Smith and Nichols, 2009] and is particularly suitable for small sample sizes. The thresholded group-level maps were anatomically labeled using the bspmview toolbox (www.bobspunt.com/software/bspmview/; Spunt, 2016).

##### Changes in regFC across transfer runs

We applied a psychophysiological interaction (PPI) analysis to investigate changes in FC between target SAN/DMN ROIs and the whole brain, modulated by task blocks during transfer runs [McLaren et al., 2012; O’Reilly et al., 2012], using the toolbox CONN (version 19.c) [Whitfield-Gabrieli and Nieto-Castanon, 2012]. Seed-based PPI maps were estimated across pre-training, post-training, and follow-up sessions using a two-level analysis. As seeds we defined the four ROIs that comprised the SAN and the four DMN ROIs that were targeted during neurofeedback training. The SAN and DMN regions were masked with subject-specific gray matter maps prior to their time-course extraction. For the first-level analysis, the interactions between the task blocks and the time courses of the targeted regions were defined as regressors of interest in separate GLMs for each seed and the betas were estimated. Regressors of no-interest were defined as the six realignment parameters, their first-level derivatives, and the five principal components from white matter and CSF time-series (Behzadi, Restom, Liau, & Liu, 2007). Additional denoising included bandpass filtering (0.008-0.09 Hz), despiking, and linear detrending. For the second-level analysis, beta images of all participants were entered into Wilks’ Lambda tests (a multivariate approach alternative to the repeated-measures ANOVA, robust against the violation of the compound-symmetry assumption). The group variance was then inferred across pre-training, post-training, and follow-up sessions for each seed. Thresholded statistical t-value maps were generated using the Gaussian random-field theory [Worsley et al., 1996] with a cluster-level threshold of p < 0.05, FWE (family-wise error)-corrected for multiple comparisons, and a voxel-level inclusion threshold of p < 0.001. Post-hoc analyses were performed to determine pairwise differences within the resulting PPI clusters across sessions using the library ‘emmeans’ in RStudio with p < 0.05, Tukey-corrected for multiple comparisons. Brain areas where regFC changes were found were anatomically labeled using xjView (www.alivelearn.net/xjview).

#### 2.4.2. Resting-state functional connectivity analyses

##### Seed-based rsFC

We used rsFC to investigate changes in FC between target ROIs and the whole brain at rest due to neurofeedback training. This rsFC analysis was performed using the CONN toolbox. Seed-based rsFC maps were estimated using two-level analyses across pre-training, post-training, and follow-up sessions. As seeds we defined the four ROIs that comprised the SAN and the four DMN ROIs that were targeted during neurofeedback training, masked with subject-specific gray matter maps prior to their time-course extraction. For the first-level analysis, the seed-based time-courses were defined as regressors and beta values were estimated voxel-wise for each participant and region using GLMs. The regressors of no-interest included the six realignment parameters and their first-level derivatives, and the five principal components from white matter and CSF time-series. Denoising included bandpass filtering, despiking, and linear detrending. For the second-level analysis, beta images of all participants were entered into a Wilks’ Lambda test and the group variance was inferred across sessions. Thresholded statistical t-value maps were generated using Gaussian random-field theory with a cluster-level threshold of p < 0.05, FWE-corrected for multiple comparisons, and a voxel-level inclusion threshold of p < 0.001. Post-hoc analyses were performed to determine pairwise differences across sessions within the thresholded clusters. Brain areas where rsFC changes were found were anatomically labeled using xjView.

##### Changes in rsFC

We investigated modulations in rsFC within the SAN and DMN ROIs across pre-training, post-training, and follow-up sessions using a graph theoretical approach. In graph theory applied to neuroimaging, the degree of FC is defined as the number of edges of an individual node for a given network and a given threshold [Rubinov and Sporns, 2010]. Here, the degree of FC estimates to which extent a target network region is connected to the rest of the brain. We computed the degree of rsFC using the intrinsic connectivity distribution (ICD) approach [Scheinost et al., 2012], which does not require the choice of an arbitrary threshold. Specifically, for this analysis, slice-time corrected and realigned resting-state functional images were first normalized and resampled to a voxel size of 4×4×4 mm^3^, to reduce computational load in ICD computation, and smoothed using a kernel of 8 mm FWHM. The ICD was computed voxel-wise and for each participant and session using a customized code as reported in [Scheinost et al., 2012]. To assess changes in the degree of SAN and DMN regions, we averaged the ICD voxel values within these regions for each participant and session. One-way repeated-measures ANOVAs were computed for each region, with session being defined as within-subject factor. Post-hoc analyses were performed to determine pairwise differences across sessions. p-values were adjusted for multiple comparisons at the region level using the Tukey method. We estimated effect sizes for the main effect and the pairwise comparisons, i.e., partial η^2^ and Cohen’s d, respectively.

#### 2.4.3. Analysis of behavioral effects

We investigated changes in sustained attention across sessions, as measured by PVT. We previously reported that sustained attention improved right after neurofeedback training [Pamplona et al., 2020a]. Specifically, participants improved in the first few minutes of the PVT task, but this improvement was no longer found in later trials of the PVT. Here, we tested whether this initial improvement persisted in follow-up sessions. We used linear mixed models to account for the hierarchical structure (multiple measurements of response time for each subject), with the factors Session and Trial; Trial being a continuous variable. Since we were interested in differences in reaction time over trials across sessions, we checked whether the two-way interaction Session x Trial was significant. We then performed post-hoc analyses to pairwise compare the reaction time across sessions at early and late trials separately. The post-hoc analysis of Trial as a continuous variable was performed following the procedures described in [Cohen and Cohen, 1983; West et al., 1996]; i.e., early and late trials were defined as the average trial minus and plus one standard deviation, respectively. Subject was defined as a random factor, Session and Trial were defined as fixed factors. For linear mixed model and post-hoc analyses, we used the libraries ‘lme4’ and ‘emmeans’ in RStudio (adjusted p-values for multiple comparisons using the Tukey method), respectively. Effect sizes for post-hoc analysis following linear mixed models were estimated with the library ‘emmeans’.

In addition, we investigated changes in self-reported attention, namely those DSSQ sub-scores that were thought to be modulated between follow-up and pre-training sessions (i.e., motivation, self-focused attention, concentration, control and confidence, task-related interference). The other DSSQ score are not specific to relevant attention measures and were not tested. Also the CFQ scores were not tested here because they are assumed to be stabile over long periods [Broadbent et al., 1982]. We used paired t-tests and dependent two-group Wilcoxon signed-rank tests for parametric and nonparametric distributions, respectively, as assessed by Shapiro-Wilk tests. For each analysis, the p-values were adjusted for multiple comparisons using the false discovery rate (FDR). Furthermore, we semantically compared and described the most reported strategies for both regulation and baseline blocks, as well as how many participants kept the same strategy in the follow-up transfer runs compared to the post-training transfer runs. We also separated the participants in two groups, one comprised of participants that reported using the same strategies in both post-training and follow-up transfer runs and one that reported different strategies, and compared the betas of regulation performance between groups with a two-sample t-test. In addition, we compared the self-rated concentration level between pre-training and follow-up transfer runs with a paired t-test.

Finally, we performed an exploratory analysis in which we investigated associations of improved sustained attention with changes in functional connectivity between the DMN ROIs and the occipital gyrus. We first computed the absolute changes (i.e., the simple difference) of the average reaction time during the first half of the PVT for post-minus pre-training sessions and for follow-up minus pre-training sessions. Only the first half of the PVT was considered here since we observed attentional improvement after neurofeedback training only during the first minutes of its application. We then computed the absolute changes of regFC estimates between the occipital gyrus and the PCC, the L Ang, and the R Ang, as well the absolute change of rsFC estimate between the occipital gyrus and the R Ang, for post-minus pre-training sessions and for follow up minus pre-training sessions. These regFC and rsFC estimates were selected because of the significant findings between DMN ROIs and the occipital gyrus (Figs. 4 and 5). Finally, we computed the Spearman correlation between PVT reaction time and functional connectivity estimates separately for absolute changes post-minus pre-training sessions and follow-up minus pre-training sessions. The p-values were adjusted using the FDR for the multiple comparisons post-minus pre-training sessions and follow-up minus pre-training sessions, separately.

**Figure 2.**
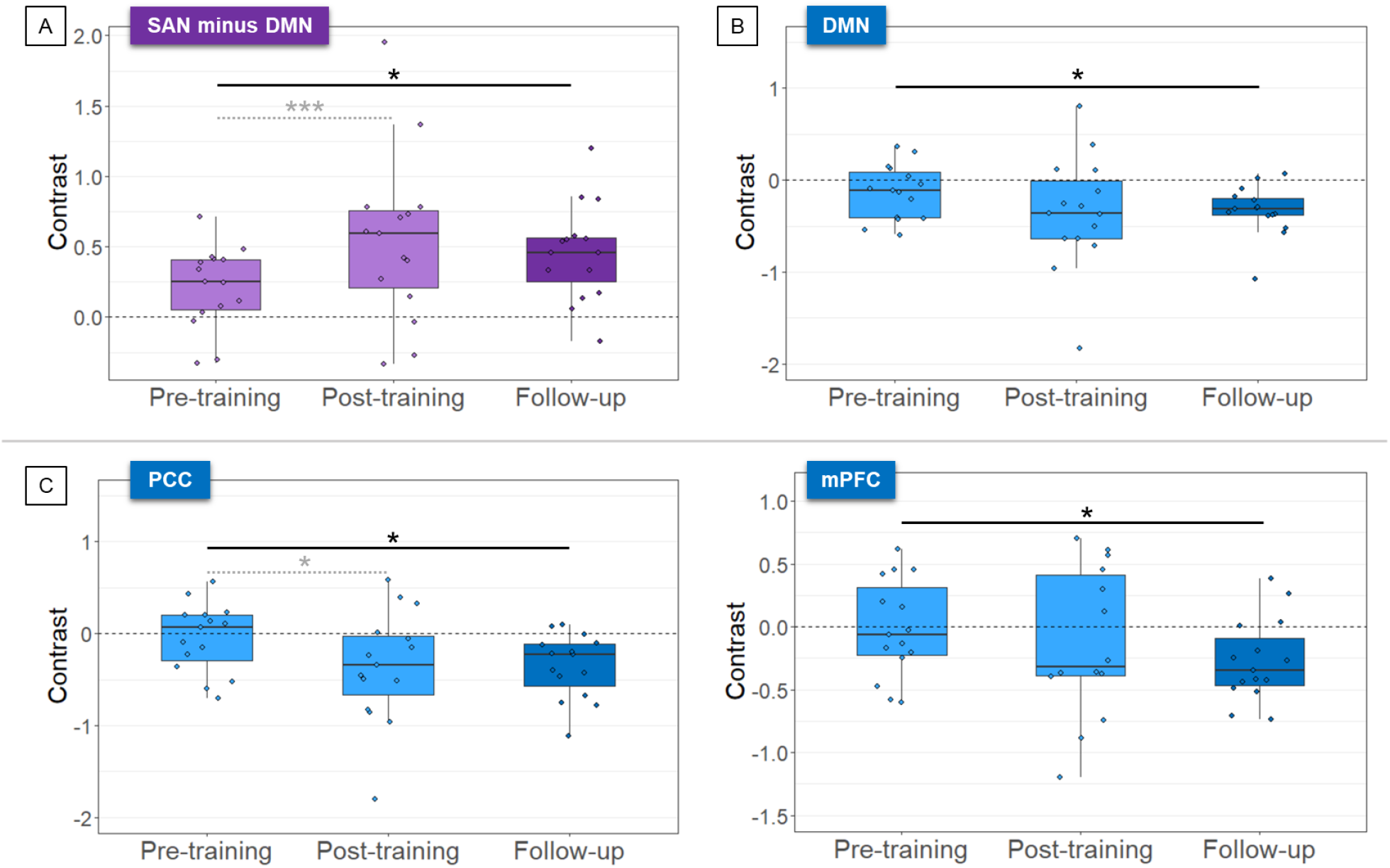
Learned self-regulation of the differential SAN-DMN activity was maintained during follow-up transfer runs two months after neurofeedback training (A). Self-regulation was mainly driven by down-regulation of the DMN (B). Specifically, the posterior cingulate cortex (PCC) and the medial prefrontal cortex (mPFC) as part of the DMN showed maintained down-regulation during follow-up runs (C). The graphs show the activation contrast between regulation and baseline blocks for pre-training, post-training, and the two follow-up transfer runs. Purple and blue colors represent the differential SAN-DMN activity and DMN regions, respectively. Light and dark colors represent pre-/post-training and follow-up sessions, respectively. The gray dashed lines represent significant differences previously reported in [Pamplona et al., 2020a]. Asterisks indicate significant session differences (*** p < 0.001, * p < 0.05, uncorrected).

**Figure 3.**
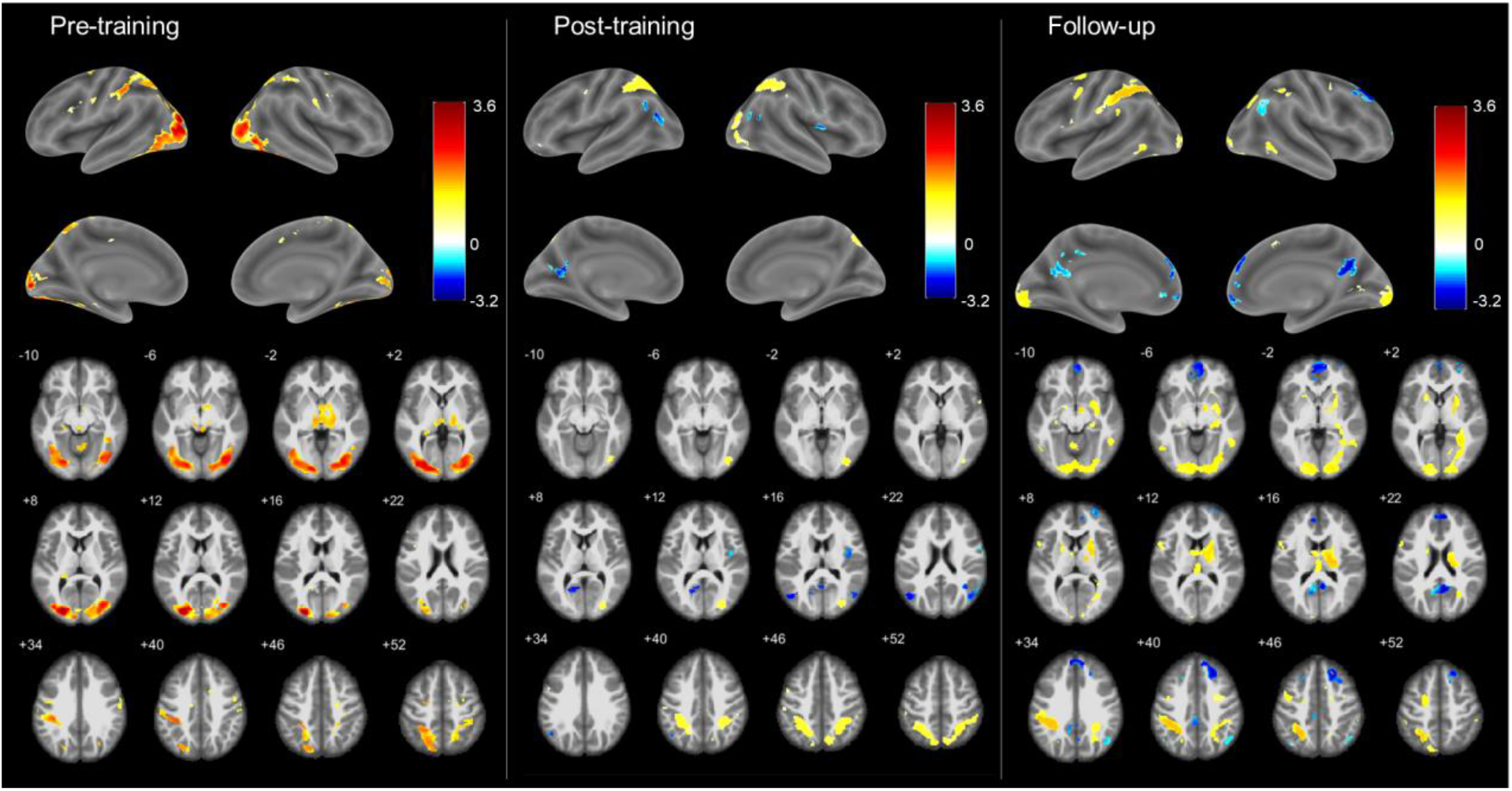
Whole-brain analyses showed greater down-regulation of the DMN during follow-up transfer sessions compared to the pre- and post-training transfer sessions. The dorsal attention network (DAN) was activated in all sessions. Left, middle, and right columns show pre-training, post-training, and follow-up transfer sessions, respectively. Hot and cold colors represent significant activation and deactivations during regulation compared to baseline blocks, respectively, overlaid on surface-rendered (top) and axial slices (bottom) from a brain template. T-maps were generated by TFCE, thresholded at p < 0.001 unc. for illustration.

**Figure 4.**
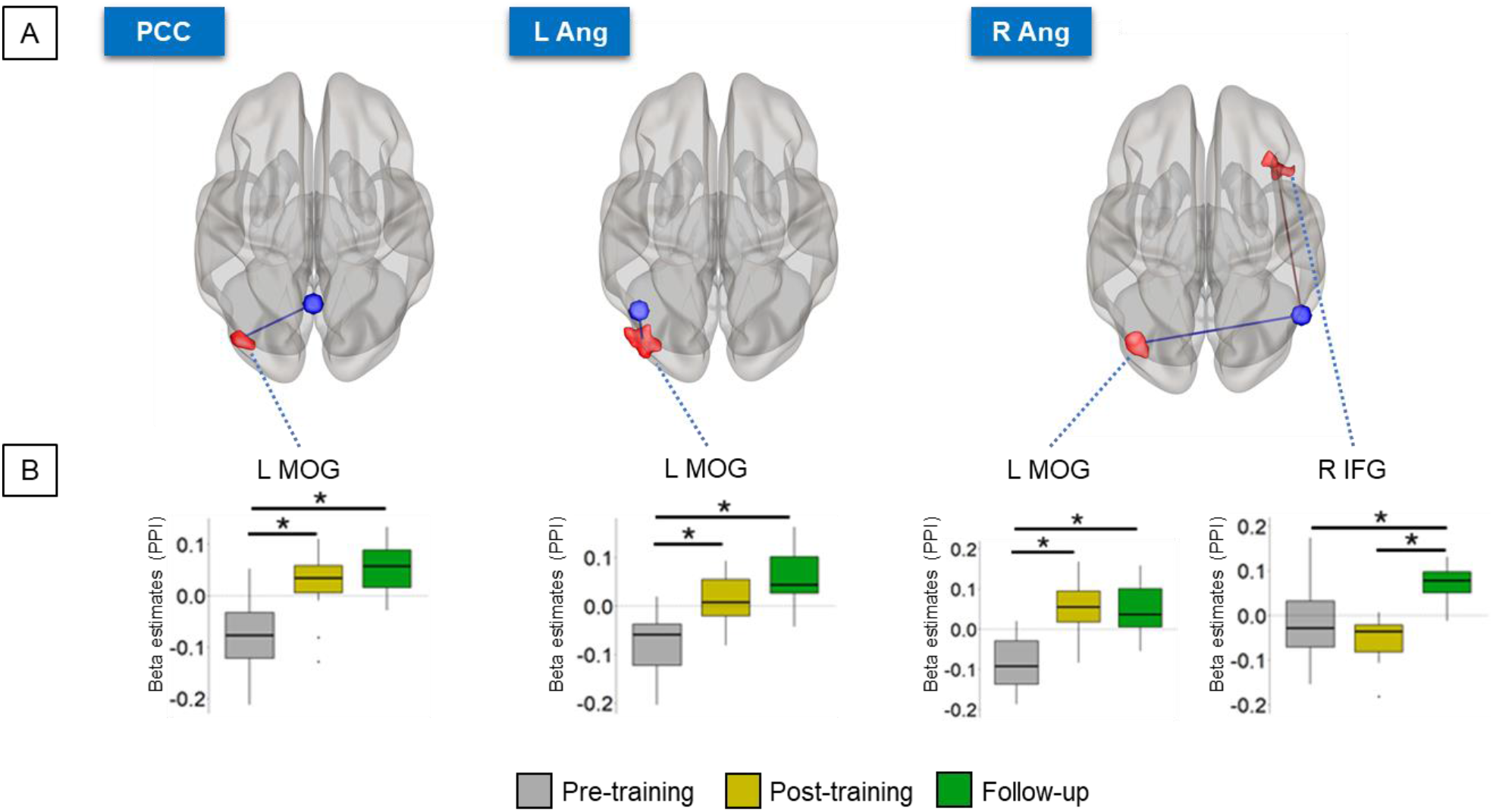
(A) Regulation-specific FC analysis showed increased FC between DMN regions (PCC, L Ang, R Ang) and the left middle occipital gyrus during post-training and follow-up transfer runs compared to pre-training runs. Blue and red regions represent DMN ROIs and significant seed-to-voxel FC regions, respectively, projected onto glass brains. (B) Boxplots represent the individual betas estimated for the PPI regressor of the DMN ROIs for each session; gray, yellow, and green represent pre-training, post-training, and follow-up transfer sessions, respectively. Asterisks indicate significant differences corrected for multiple comparisons using the Tukey method (p < 0.05). MOG = middle occipital gyrus, IFG = inferior frontal gyrus, L/R = left/right.

**Figure 5.**
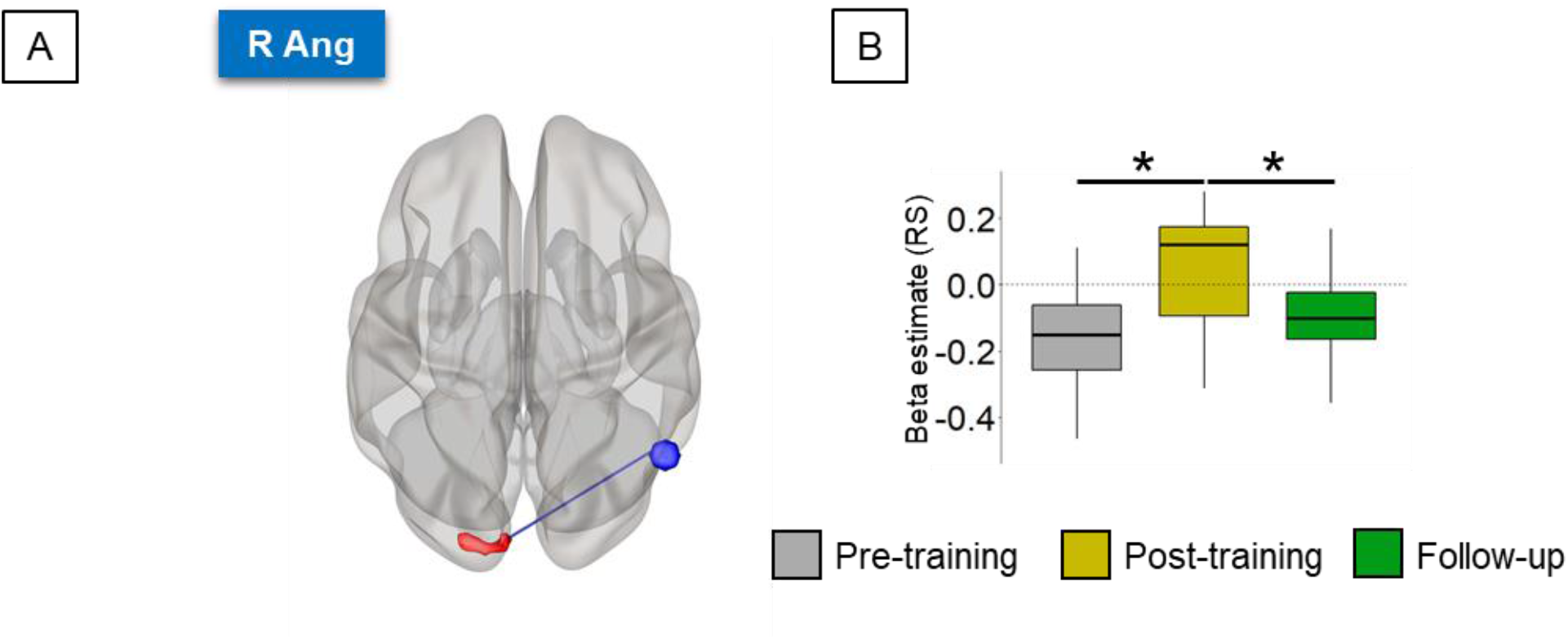
(A) Resting-state (RS) FC between the R Ang DMN region and the middle occipital gyrus increased during post-training compared to pre-training runs, but returned to pre-training levels during the follow-up runs. Blue and red brain areas represent DMN regions and significant seed-to-voxel rsFC regions, respectively, projected onto a glass brain. (B) Boxplots represent the individual betas estimated for the regressor constructed with the average time-course within the R Ang; gray, yellow, and green represent pre-training, post-training, and follow-up sessions, respectively. The dashed black lines in the boxplots represent the zero level. Asterisks indicate significant differences corrected for multiple comparisons using the Tukey method (p < 0.05). R Ang = right angular gyrus.

## 3. RESULTS

### 3.1 Long-term effects of neurofeedback training during follow-up transfer runs

#### 3.1.1 Long-term effects in the trained networks during follow-up transfer runs

Previously we found that participants learned to control the differential activity SAN-DMN, mainly through down-regulating the DMN [Pamplona et al., 2020a]. Here, our new results show that learned self-regulation of the differential SAN-DMN activity was maintained during transfer runs without feedback two months after the neurofeedback training (paired t-test between pre-training and follow-up runs: t(14) = 1.92, d = 0.51, p = 0.038 (Fig. 2A). Also during follow-up runs, self-regulation was primarily driven by a persistent down-regulation of the DMN (paired t-test between pre-training and follow-up runs: t(14) = -1.80, d = -0.46, p = 0.047 (Fig. 2B). Compared to pre-training, self-regulation of the SAN activity was neither different during post-training runs nor during follow-up runs (paired t-test between pre-training and follow-up runs: t(14) = 0.59, d = 0.15, p = 0.28) (Fig. S1A).

When analyzing self-regulation performance of each of the SAN and DMN ROIs separately, we observed that increased ability in down-regulating the PCC was maintained during follow-up runs (t(14) = 2.49, d = -0.64, p = 0.013) (Fig. 2C). Furthermore, the mPFC was down-regulated during follow-up (t(14) = -2.94, d = -0.50, p = 0.037) (Fig. 2C), but not during the post-training session. No other regulation effects within SAN / DMN ROIs changed significantly across transfer runs (Fig. S1B).

#### 3.1.2 Long-term effects during follow-up transfer runs across the whole brain

Whole-brain analyses showed significant deactivation (i.e., estimated betas at regulation < baseline) in the DMN during the follow-up transfer session (i.e., averaged over the two follow-up runs) (Fig. 3 and Table 1). Brain areas showing deactivation in the PCC and mPFC were larger in the follow-up compared to the post-training session. While the right IPS was activated during all transfer runs and the bilateral angular gyri were deactivated during the post-training run, all DMN ROIs were deactivated during follow-up transfer runs. Activation in the dorsal attention network (DAN) was detected in all transfer sessions. The thalamus was also activated in the follow-up session. A complete list of activated and deactivated brain areas is reported in Table 1. The contrasts post- vs. pre-training and follow-up vs. pre-training showed decreased activity in the left and right middle occipital gyrus, respectively (Fig. S2 and Table 1).

**Table 1.**
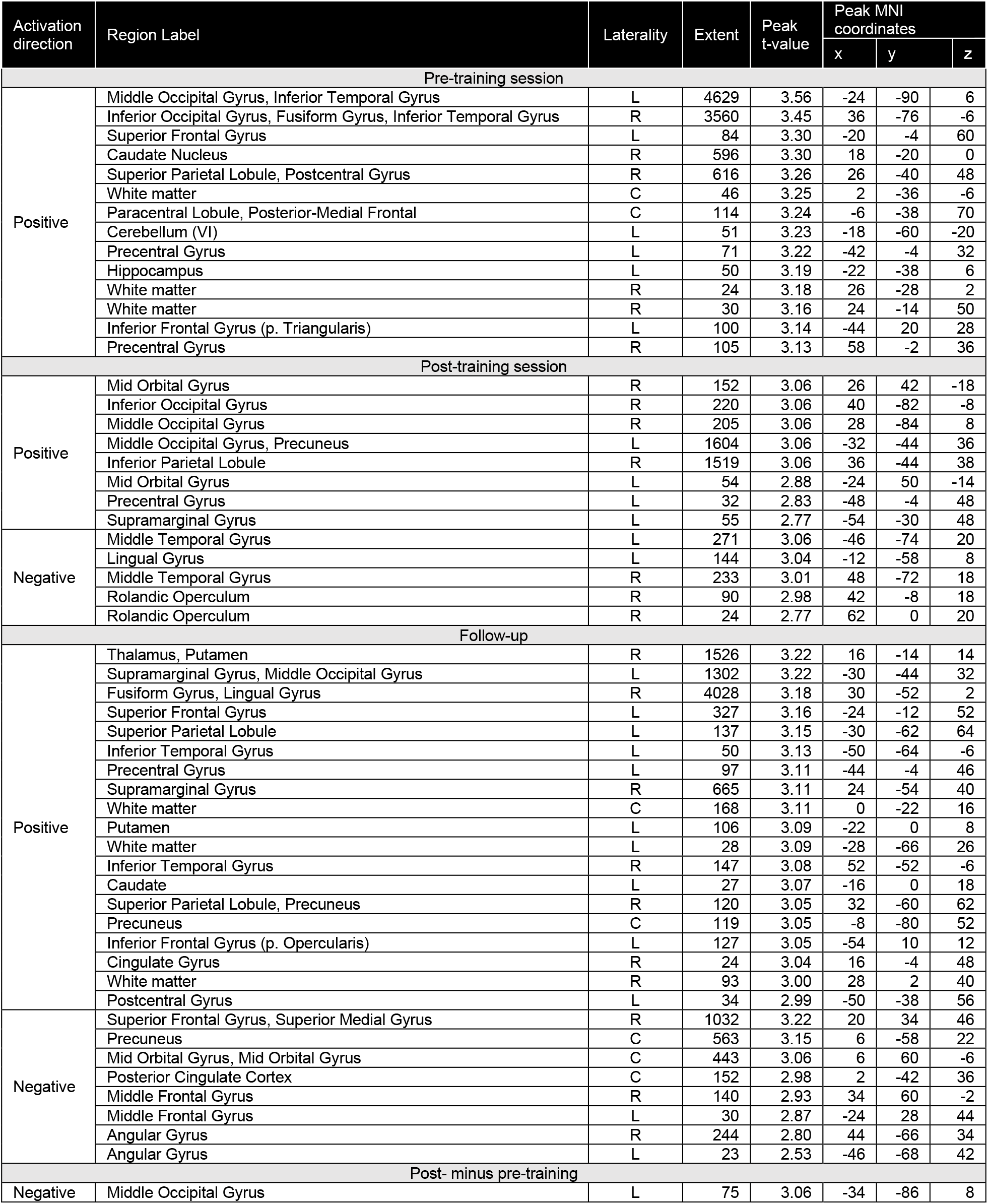
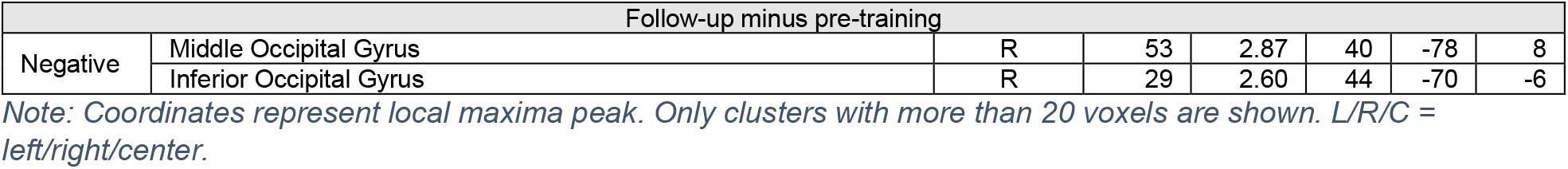
Significant positive and negative activations during the pre-training, post-training, and follow-up transfer sessions, as shown in Figure 3, including contrasts for the post-minus pre-training and follow-up minus pre-training sessions, as shown in Figure S2.

#### 3.1.3 Regulation-specific FC changes across transfer runs

Significant regFC changes between pre-training, post-training, and follow-up transfer runs were found mainly between the DMN regions PCC, L Ang, R Ang and the left middle occipital gyrus (Fig. 4). RegFC changes were also found between the SAN regions and the right angular gyrus, left hippocampus, and postcentral gyrus (Fig. S3). Summary group results of brain areas with significant regFC changes are shown in Table 2.

**Table 2.** Changes in regulation-specific (regFC) and resting-state (rsFC) functional connectivity between SAN/DMN and other brain areas across sessions, as illustrated in Figs. S3, 4, and 5.

| regFC |  |  |  |  |  |  |  |  |  |  |  |  |
| --- | --- | --- | --- | --- | --- | --- | --- | --- | --- | --- | --- | --- |
| Seed | Connecting brain area |  | Extent (voxels) | Follow-up minus pre-training |  |  | Post-training minus pre-training |  |  | Follow-up minus post-training |  |  |
|  | Label | MNI peak coordinates (mm) |  | t(28) | Cohen's d | p-value | t(28) | Cohen's d | p-value | t(28) | Cohen's d | p-value |
| SAN1 | Right angular gyrus | (34, -66, 44) | 120 | +5.38 | +2.16 | <0.0001 | - | - | - | +3.97 | +1.29 | 0.0013 |
| SAN2 | Left postcentral gyrus | (-44, -30, 54) | 142 | -4.13 | -1.41 | 0.0008 | - | - | - | -5.73 | -2.29 | <0.0001 |
| SAN3 | Left hippocampus | (-18, -8, -20) | 133 | +4.86 | +2.00 | 0.0001 | - | - | - | +4.66 | +1.68 | 0.0002 |
| SAN4 | Left postcentral gyrus | (-44, -16, 48) | 125 | -3.16 | -1.31 | 0.010 | +2.50 | +1.00 | 0.05 | -5.66 | -2.46 | 0.0001 |
| DMN1 | Left middle occipital gyrus | (-42, -82, 12) | 121 | +4.20 | +1.52 | 0.0007 | +5.44 | +2.14 | <0.0001 | - | - | - |
| DMN3 | Left middle occipital gyrus | (-46, -82, 8) | 165 | +5.83 | +2.09 | <0.0001 | +3.97 | +1.53 | 0.0013 | - | - | - |
| DMN4 | Left middle occipital gyrus | (-40, -76, 10) | 113 | +4.79 | +2.09 | <0.0001 | +4.79 | +2.08 | <0.0001 | - | - | - |
|  | Right inferior frontal gyrus | (36, 24, -16) | 136 | +4.11 | +1.45 | 0.0009 | - | - | - | +5.76 | +2.69 | <0.0001 |
| rsFC |  |  |  |  |  |  |  |  |  |  |  |  |
| DMN4 | Left superior occipital gyrus | (-6, -94, 18) | 87 | - | - | - | +5.35 | +1.23 | <0.0001 | -3.67 | -0.87 | 0.0028 |

*Note: Results from post-hoc analyses of the pairwise differences (follow-up minus pre-training, post-training minus* *pre-training, and follow-up minus post-training). Blank cells indicate non-significant contrasts.*

### 3.2 Changes in seed-based rsFC

Significant seed-based rsFC changes across pre-training, post-training, and follow-up resting-state sessions were found between the right angular gyrus and the superior occipital gyrus (Fig. 5). Summary group results of the regions with significant rsFC changes is shown in Table 2.

Graph-theoretical analysis revealed that the degree of rsFC changed over the course of pre-training, post-training, and follow-up resting-state sessions in two of the trained regions: right IPS (F(2,28) = 4.40, η^2^ = 0.24, p = 0.022) and PCC (F(2,28) = 3.86, η^2^ = 0.22, p = 0.03) (Fig. 6). Post-hoc analyses showed that the degree of rsFC increased in the right IPS from pre-training to follow-up (mean_pre-training_ = 0.08 ± 0.02, mean_follow-up_ = 0.10 ± 0.03, t(28) = 2.41, d = 0.93, p = 0.03) and in the PCC from post-training to follow-up (mean_post-training_ = 0.07 ± 0.02, mean_follow-up_ = 0.09 ± 0.02, t(28) = 2.78, d = 0.79, p = 0.03).

**Figure 6.**
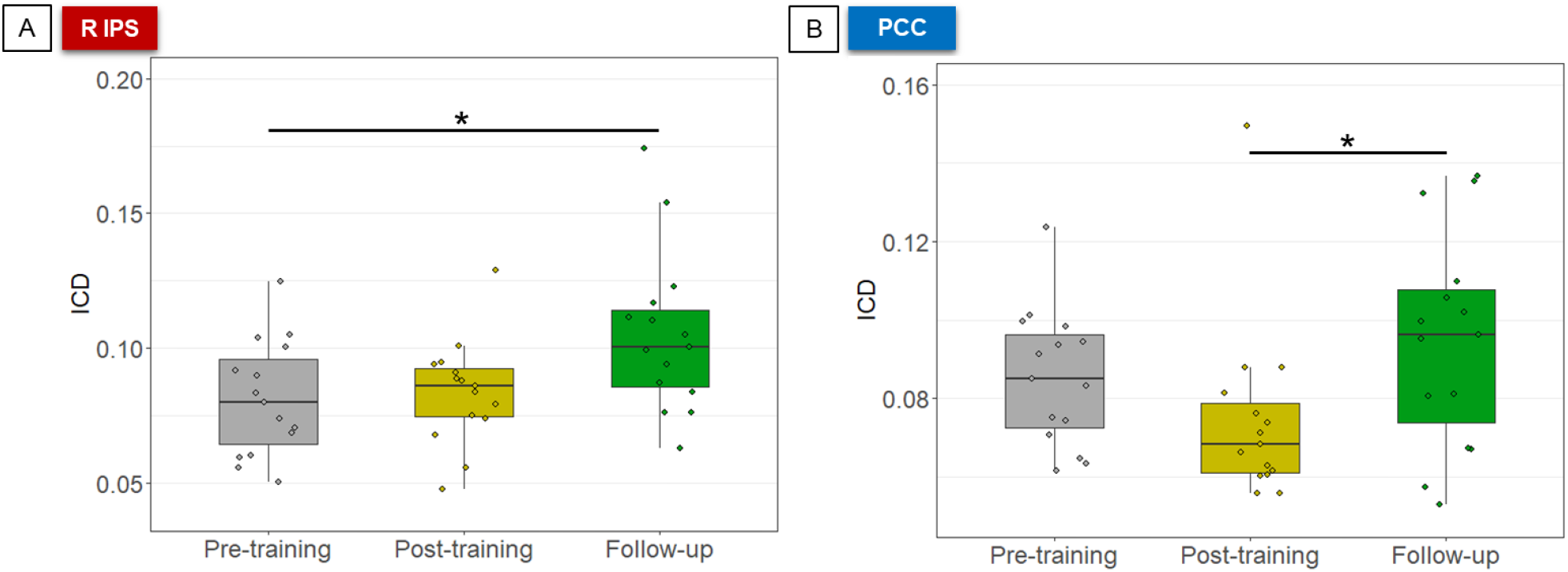
Higher degree of rsFC was observed in the follow-up session in (A) the right IPS (compared to pre-training), and (B) the PCC (compared to post-training). Asterisks indicate significant differences in post-hoc analyses, corrected for multiple comparisons using the Tukey method (p < 0.05). R IPS = right intraparietal sulcus, PCC = posterior cingulate cortex.

### 3.3 Behavioral effects cease to exist

We previously found that neurofeedback training led to shorter reaction times in early trials of the PVT (effect size = 0.15), indicating improved sustained attention in the first minutes of the task following neurofeedback training [Pamplona et al., 2020a]. However, the training-induced improved sustained attention in early trials of the PVT was not maintained in follow-up tests two months after the training (Fig. 7). Specifically, there was a significant interaction between the factors Day and Trial (F(1,5091) = 6.58, p = 0.0014). According to the procedure described for post-hoc analysis following linear mixed models with continuous variables [Cohen and Cohen, 1983; West et al., 1996], reaction time for the PVT during early trials in the follow-up was longer than in the post-training session (t(5091) = 2.40, effect size = 0.12, p = 0.04; follow-up: M = 343 ms, CI = [318, 368] ms; post-training: M = 336 ms, CI = [311, 361] ms) and was not different from the pre-training session (t(5091) = 0.66, effect size = 0.03, p = 0.8; pre-training: M = 344 ms, CI = [320, 369] ms). In addition, the PVT reaction time during late trials was longer in the follow-up compared to the pre-training session (t(5091) = 3.60, effect size = 0.18, p = 0.0010; follow-up: M = 361 ms, CI = [336, 386] ms; pre-training: M = 351 ms, CI = [326, 376] ms), and was not different from the post-training session (t(5091) = 2.07, effect size = 0.10, p = 0.10; M = 356 ms, CI = [331, 381] ms).

**Figure 7.**
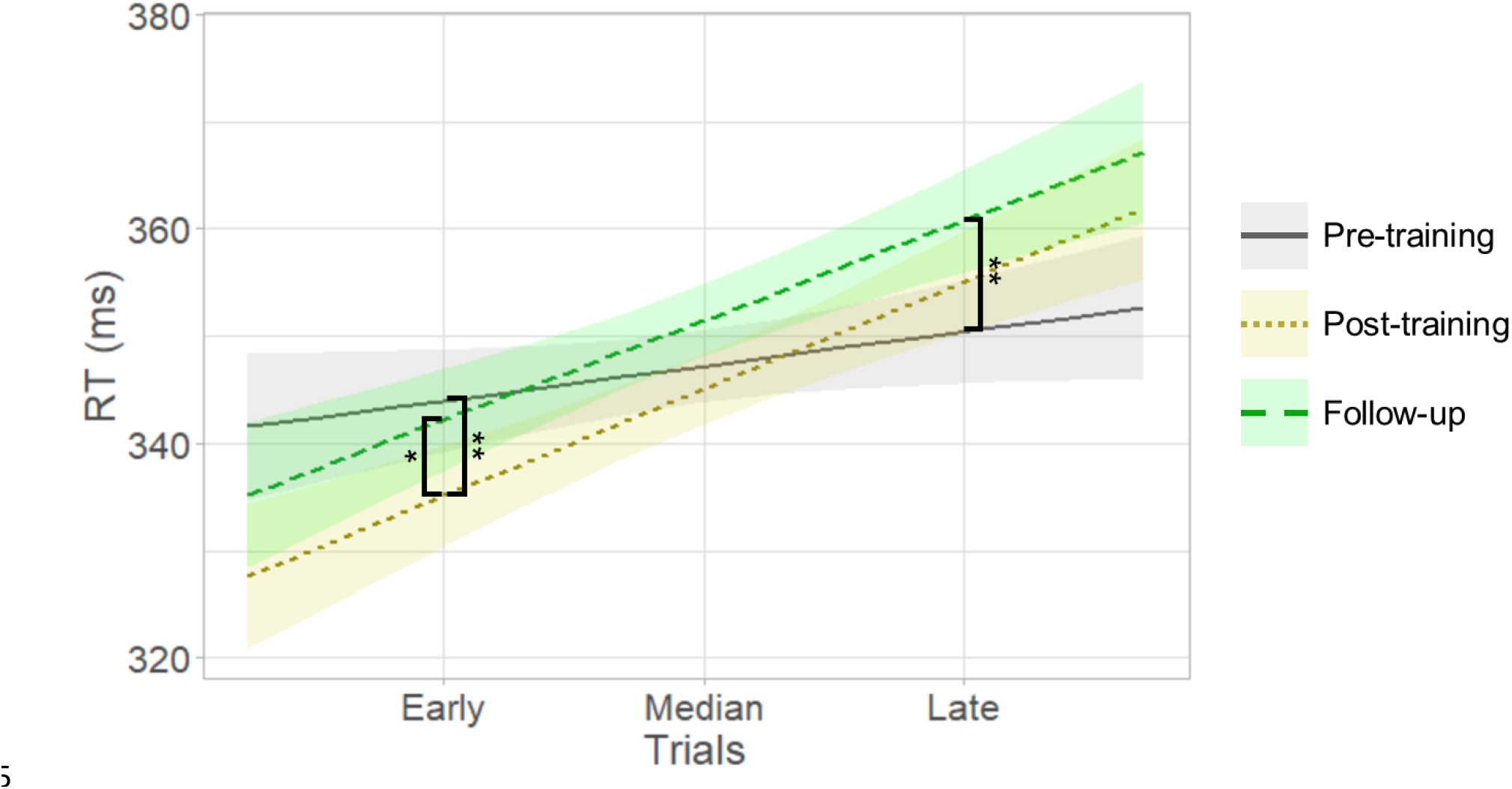
Differences between PVT reaction time (RT) in pre-, post-training, and follow-up sessions indicate that improved sustained attention after neurofeedback training was no longer evident two months later. Also, during follow-up, performance during late trials was worse compared to pre-training. Gray, yellow, and green colors represent measurements at pre-training, post-training, and follow-up sessions, respectively. Asterisks indicate significant differences in post-hoc analyses, corrected for multiple comparisons using the Tukey correction (** p < 0.01, * p < 0.05).

Attentional/motivational states, measured by the DSSQ, during follow-up transfer runs were not different from pre-training sessions (all FDR-corr. ps > 0.05). A list of strategies used for regulation and baseline blocks during pre-training, post-training, and follow-up transfer sessions is shown in Table S2. The most reported strategies for regulation blocks were keeping the attentional focus on the geometry of the up-arrow (N = 6), thoughts related with past memories or future projection (2), and performing mental math (2). The most reported strategies for baseline blocks were trying to think about nothing in particular (4), mind-wandering (3), and mental imagery of sports (2). Eight of the participants reported the same strategies (for both blocks) that they reported for transfer runs right after the end of the training. Considering strategies adopted in the follow-up and the post-training run, self-regulation performance was not different between participants that used the same reported strategies and participants that used different ones (Welch two sample t-test: T(7.4) = 0.07, p = 0.9). There were no differences between self-rated concentration reported after pre-training and follow-up transfer runs (paired t-test: T(12) = 0.97, p = 0.4).

In an exploratory analysis, we found that the absolute change in the regFC between the L Ang/R Ang and the occipital gyrus was correlated with the absolute change in the reaction time in the first half of the PVT across individuals and considering the difference post-minus pre-training runs (L Ang: r = -0.61, p = 0.04; R Ang: r = -0.66, p = 0.04). The significant negative correlation indicates that the degree of regFC increase during the post-training run is associated with the degree of reaction time reduction. The reaction time change in the first half of the PVT was not correlated with the post-minus pre-training change in the regFC between the PCC and the occipital gyrus (r = -0.23, p = 0.5), nor with the post-mins pre-training change in the rsFC between the R Ang and the occipital gyrus (r = -0.12, p = 0.7). Correlations considering follow-up minus pre-training changes were not significant (all ps > 0.05).

## 4. DISCUSSION

In this study, we investigated the maintenance of brain and behavioral changes associated with network-based fMRI neurofeedback training for sustained attention during transfer and resting-state runs conducted before, one day after, and two months after the training. We found evidence for maintenance of learned self-regulation and lasting, plastic brain changes. Specifically, we found that after two months, participants were still able to up-regulate the differential SAN-DMN activity, and that successful self-regulation was driven mainly by down-regulating the DMN. Also, the training-induced increase in functional coupling between DMN and occipital cortex during transfer runs was maintained. Finally, the degree of FC during resting-state runs increased in those brain regions that were successfully trained with neurofeedback. On the other hand, the increase in functional coupling between DMN and occipital cortex during resting-state runs after training returned to baseline level during follow-up runs. Behaviorally, the improved sustained attention right after neurofeedback training also returned to baseline level two months later.

### 4.1 Lasting neurofeedback effects on the differential SAN-DMN activity

The ability to self-regulate differential SAN-DMN activity, acquired through neurofeedback training, was still present two months after training. This is in line with previous findings showing that brain self-regulation learned through neurofeedback training is maintained for months [Amano et al., 2016; Robineau et al., 2017]. Whereas these previous studies trained for three sessions, neurofeedback training in our study was limited to two sessions of 45 min each, showing that relatively short neurofeedback training allows participants to learn lasting self-regulation skills (i.e., at least two months).

Learned self-regulation of the differential feedback signal was primarily driven by down-regulation of the DMN, which was observed right after training and during follow-up two months later(Figs. 2 and 3). Interestingly, DMN down-regulation during transfer runs was even more pronounced during follow-up compared to right after training. For example, down-regulation of the mPFC and right angular gyrus was observed only during follow-up after two months but not directly after neurofeedback training (Figs. 2C and 3). Akin to improvements following behavioral interventions, it might be that after training participants continue practicing self-regulation in everyday life, thus further improving [Rance et al., 2018]. Since activation in the DMN is associated with internally oriented attention [Bonnelle et al., 2011; Gusnard et al., 2001; Hinds et al., 2013; Mason et al., 2007], the improvement in DMN down-regulation over time might reflect a reduced propensity for mind-wandering in favor of a greater externally oriented attention. Therefore, learning and maintenance of self-regulation of large-scale networks might have implications in the ability of censoring spontaneous task-irrelevant thoughts. However, these remain speculations as we currently do not have data on practice outside the experiment and reduced mind-wandering.

Apart from DMN down-regulation, the only SAN ROI that was up-regulated in the post-training session was the right IPS. However, this was not maintained in the follow-up session (Fig. S1). The IPS is part of the DAN, which was active during all transfer runs (Fig. 3). DAN activity is related to the preparation and execution of top-down or goal-directed attention (Fox et al., 2005) and the IPS, specifically, is associated with reorienting top-down attention [Corbetta and Shulman, 2002]. At least 40% of the participants used a strategy that involves reorienting top-down attention (“constantly reorienting the attentional focus on the geometry of the up-arrow” (Table S2), thus likely resulting in activation of the DAN (Fig. 3). The increased ability to activate IPS during the post-compared to the pre-training session might indicate improved engagement of neural resources mediating top-down attention. However, the follow-up session showed that the lasting effects were unrelated to the SAN, but primarily driven by lasting increased DMN down-regulation. The lack of lasting increased SAN up-regulation might have been a consequence of having trained healthy participants with intact top-down attention. It remains to be tested if our neurofeedback training in patients with attention-deficits would lead to lasting SAN (and DMN) changes.

When comparing post-training and follow-up transfer runs to pre-training transfer runs, we found reduced activity in the occipital gyri (Figs. 3 and S2). Hence, for participants who focused their attention on visual features of the feedback display during training (Table S2), reduced occipital activity might indicate habituation to re-occurring visual stimuli [Weigelt et al., 2008]. Alternatively, for participants who focused on internally-oriented attention (see, for example, participants #2 and #8 in Table S2), reduced occipital gyrus activity might also indicate reduced processing of visual information as a function of training sustained, internal-focused attention [Benedek et al., 2016].

Finally, we observed increased thalamus activity during follow-up transfer runs (Fig. 3). Thalamic activity is related to general arousal maintaining alertness [Sarter et al., 2001]. Thalamic activity during vigilant attention decreases over time, but returns when new conditions are presented, playing a role in compensatory attentional effort [Langner and Eickhoff, 2013]. Since we observed thalamic activity during self-regulation a long time after the end of training, it might be that the thalamus activity was associated with arousal related to compensating for a less automatic state of sustained attention, compared to directly after training.

### 4.2 Lasting neurofeedback effects on functional connectivity vs. transient behavioral changes

Our neurofeedback training induced both short- and long-term changes in regFC and rsFC (Table 2). Previous studies have reported changes in FC due to neurofeedback training in patients [Scheinost et al., 2013; Yuan et al., 2014] and healthy participants [Megumi et al., 2015; Zhang et al., 2013]. As argued by Rance and colleagues, changes in FC may be continuously reinforced after neurofeedback training [Rance et al., 2018] over the course of days [Harmelech et al., 2013], weeks [Yuan et al., 2014], or months [Megumi et al., 2015]. Our findings further support claims that neurofeedback can induce FC changes that are maintained for several months.

The most consistent FC changes that we observed were related to increased FC between the DMN and the occipital cortex (Figs. 4 and 5). More specifically, we observed that the regFC between DMN and occipital cortex increased during post-training and follow-up runs compared to pre-training runs. A previous PPI study showed that, when the frontoparietal control network is engaged, the connectivity between DMN and the occipital cortex increases [Karten et al., 2013]. It was suggested that the increase in the DMN-occipital cortex connectivity reveals a competitive interaction suppressing the bottom-up visual stream [Karten et al., 2013] and protecting internal attentive processes from potentially distracting sensory stimulation [Benedek et al., 2016]. In addition, the suppression of externally and internally distracting information, i.e., generated in the visual cortex and the DMN, respectively, is closely linked to each other and predictive of task performance [Anticevic et al., 2012]. In our study, the SAN has components from the frontoparietal control network, specifically the aMCC and the rTPJ. The aMCC was clearly engaged during the post-training and follow-up runs (Fig. S1). Therefore, the engagement of frontoparietal control network during transfer runs might have also increased the connectivity between DMN and occipital cortex. In fact, we observed that greater regFC between DMN and occipital cortex were associated with faster response time (Fig. 8), when comparing post-training with pre-training sessions. Such an association might indicate that participants learn to simultaneously suppress distracting externally and internally information and that this ability was also employed during the sustained attention task. As the task was conducted one day after the neurofeedback training, these effects might be lasting. However, these associations were not observed when comparing follow-up and pre-training sessions. Thus, while the improved regulation-related FC was maintained long term, the improved attentional performance was not. This dissociation indicates that learned brain self-regulation can be applied upon request, but does not necessarily translate in behavioral effects long-term.

**Figure 8.**
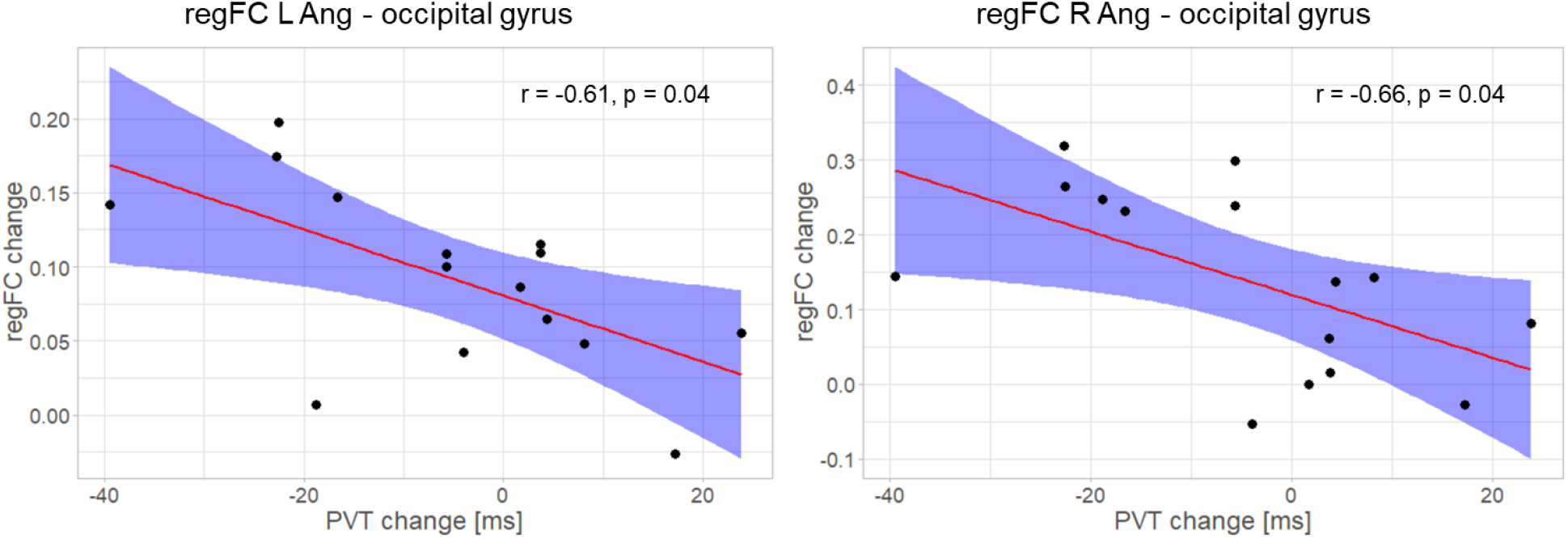
The change in the regFC between the bilateral angular gyri, brain areas that are part of the DMN, and the occipital gyrus was correlated with the reaction time change for the first half of the PVT across individuals, considering the difference post-minus pre-training sessions.

Interestingly, we also observed FC changes in resting-state runs following the end of training. First, the rsFC between DMN and occipital cortex increased one day after the end of training, but was not different from pre-training runs two months after the end of training (Fig. 5). Akin to the behavioral effects that did not last, also the rsFC changes that indicated plastic changes in the functional coupling between DMN and the occipital cortex were not maintained even though participants were still able to regulate and regFC changes persisted. However, lasting changes in the degree of FC in successfully trained ROIs during resting-state runs were observed (Fig. 6). Because during resting-state runs no active self-regulation was required, such FC changes likely represent plastic brain changes that are unrelated to concurrent mental strategies activations. And it is unlikely that they are artefactual, as the change in the degree of rsFC was specific to regions that were successfully trained with neurofeedback (Fig. 2C and Fig. S1B), probably supporting the acquired ability to regulate brain activity. Previous studies have found lasting resting-state changes following neurofeedback training [Megumi et al., 2015]. Some recent studies have even reported brain structural changes associated with neurofeedback training [Papoutsi et al., 2018; Sampaio-Baptista et al., 2021], showing the potential of neurofeedback to produce lasting effects on brain structure and processing.

Sustained attention improved to some extent right after neurofeedback training, but this improvement did not last (Fig. 7). In contrast, other studies reported persistent or even increasing behavioral effects following neurofeedback training [Amano et al., 2016; Cortese et al., 2017; Rance et al., 2018; Shibata et al., 2011]. However, also other neurofeedback studies found that behavioral effects that were present right after training did not persist. For example, an EEG-based neurofeedback study on nicotine addiction reported that short-term changes in symptom reduction were followed by a gradual return toward the baseline in the long term [Bu et al., 2019]. Why neurofeedback training sometimes induces lasting or even improving behavioral effects while sometimes such effects do not persist is a crucial question especially for clinical neurofeedback applications. Here, we can only speculate that, for example, the effect size of the initial behavioral improvement might matter. Our study trained healthy participants in a cognitive domain that we are highly trained in – attention. As a consequence, the behavioral improvement was rather small, possibly due to ceiling effects. This might be different in clinical samples (e.g., Rance et al., 2018). Therefore, studies on follow-up neurofeedback should, whenever possible, contain information about effect sizes to help elucidate this argument. Another factor might be that for behavioral effects to increase over time, frequent use of learned self-regulation in everyday situations might be important. Such practice is more likely the case in clinical populations and can be promoted by, for example, electronic diaries [Zaehringer et al., 2019]. In general, the association between neurofeedback-induced brain changes and behavioral effects remains yet to be clarified. For example, Shibata et al. (2011) found improved perceptual sensitivity after neurofeedback training even when participants did not actively self-regulate their visual cortex activity, whereas another study found that sensitivity improved only when participants actively up-regulated visual cortex activity [Scharnowski et al., 2012]. In the present study we found lasting brain changes but the behavioral effects were only transient. Only the rsFC brain changes showed the same pattern as the behavioral effects: they were present during post-training runs but no longer during follow-up runs. Following this temporal coincidence, one might speculate that rsFC changes might serve as a correlate for behavioral effects, but such a speculation requires further investigation.

Also, the relationship between mental strategies and behavioral changes requires further investigation. In our study, the individual choice of mental strategies cannot easily explain that sustained attention improved one day after neurofeedback training but no longer during follow-up 2 months later. Most subjects used the same strategies in both sessions and most of these strategies were closely related to externally- and internally-oriented focused attention during regulation and baseline blocks, respectively. It is worth mentioning that, in the follow-up session, participants were not reminded of their strategies adopted during the initial training. Further, performance during follow-up transfer runs was not dependent on whether participants used the same strategy as right after training or a different one. Thus, learned self-regulation did not seem to depend on remembering and applying the exact mental strategy that was adopted during training.

### 4.3 Outreach

First, whenever possible, one should include follow-up neuroimaging assessments of functional and/or anatomical plasticity due to the training, rather than only shortly after intervention or only behavioral or regulation-specific measurements. Although still specific to the MR setting, resting-state assessments may better reflect transfer effects of neurofeedback training because they are independent of self-regulation efforts. Follow-up neuroimaging measurements can help indicate neural reshaping over time after completed interventions [Robineau et al., 2017]. If neurofeedback-induced effects continue to increase over time, measuring them only shortly after a training intervention may lead to undervalued or undetected behavioral effects [Rance et al., 2018]. Importantly, follow-up sessions may help consolidate neuroscientific theories using neurofeedback as a causal intervention [Sulzer et al., 2013b] and define biomarkers as targets for neurotherapy [Yamada et al., 2017]. Second, since it is desired in a clinical setting that a given intervention converts practice into enduring effects, follow-up assessments can justify whether the proposed neurofeedback approach is a meaningful alternative for therapy. Therefore, neurofeedback studies that address symptoms should always rely on follow-up evaluations. Third, we note that, while clear long-lasting effects in terms of neural self-regulation may exist, persistent behavioral changes can eventually be dissociated from brain findings [Sitaram et al., 2017]. Therefore, follow-up evaluations of behavioral effects should also be conducted whenever possible. An eventual brain-behavior dissociation may raise questions about the utility of a proposed neurofeedback approach for modulating behavior or mitigating symptoms in an out-of-scanner scenario, the strategic choice of sensitive psychometric instruments, and the characterization of the targeted population. Fourth, we argue that, whenever possible, neurofeedback training and resting-state/psychometric acquisitions should be made on different days, since sleep plays an important role in consolidating learning and producing lasting changes in the brain [Walker and Stickgold, 2004]. Fifth, our study provides evidence that short sessions are sufficient (two training sessions of 45 min on separate days) to produce long-term effects [Rance et al., 2018] in terms of regulation of brain activity and connectivity changes.

### 4.4 Limitations

The main limitation of this study is that the follow-up assessments did not include a control group. While the original study included at least a control group that performed the psychometric tasks without neurofeedback training, the present analysis does not include a behavioral nor a neurofeedback control group. Therefore, we cannot conclude with certainty that the observed brain changes were caused by neurofeedback training. The brain changes could, in principle, be due to, for example, spontaneous fluctuations over time, habituation to the MR environment, or fatigue. On the other hand, the fact that the post-training self-regulation results were reproducible during follow-up runs two months later and the fact that the brain changes were predominantly specific to the trained brain areas suggest that the brain changes were indeed associated with neurofeedback training.

A second major limitation is the modest sample size. Resource constraints like limited MR scanner availability and scanning costs make scanning larger samples difficult, especially because participants in neurofeedback experiments are being scanned repeatedly. With N = 15 and each of these subjects having been scanned on 5 different days (resulting in a total of 75 MR acquisitions), this study is well within the standard range for fMRI-based neurofeedback studies [Fede et al., 2020]. To accommodate the moderate sample size statistically, non-parametric tests such as TFCE for statistical mapping in low sample sizes were used.

Finally, the follow-up session was acquired two months after the end of the training. While two months seem sufficiently long to assess lasting effects that go beyond immediate post-training changes, other studies showed that neurofeedback training effects can last much longer [Amano et al., 2016; Ramot et al., 2017; Robineau et al., 2017; Zilverstand et al., 2015b]. Hence, from one follow-up after two months we cannot infer the temporal course and an upper bound for neurofeedback training effects.

## 5. CONCLUSION

The goal of neurofeedback training is to modulate behavior, emotion, cognition, or clinical symptoms long-term through self-regulating brain activity. To evaluate whether this ambition has been achieved, follow-up assessments are key. We found that two months after the end of neurofeedback training participants were still able to exert self-regulation of the differential SAN-DMN activity, and this during transfer runs without feedback. Lasting brain changes also included FC measures of the trained ROIs to other brain regions in runs during which participants engaged in active self-regulation as well as during resting-state runs without concomitant self-regulation. These results provide information on important facets of follow-up assessments: (a) maintenance of the initially learned self-regulation skill (i.e., SAN-DMN regulation), (b) maintenance of brain changes related to self-regulation that go beyond the trained ROIs (i.e., FC changes during transfer runs), and (c) plastic brain changes in the absence of ongoing self-regulation (i.e., resting-state changes). Another important aspect of follow-up assessments is (d) behavioral effects. While others found behavioral effects to increase after neurofeedback training [Rance et al., 2018], the (relatively weak) behavioral effects we observed right after the training did not persist. Such a discrepancy between lasting brain changes but transient behavioral effects poses important questions regarding the brain-behavior associations above and beyond neurofeedback. Overall, this study highlights the importance of follow-up investigations of neural and behavioral changes associated with neurofeedback training, so that this promising approach can develop its full potential as a scientific and clinical tool.

## Acknowledgements

We thank Prof. Dustin Scheinost for providing technical advice on the intrinsic connectivity distribution, Dr. Alfonso Nieto-Castanon for support with CONN software, Dr. Philipp Stämpfli for technical support with imaging, and Prof. Christian Gaser for advice on the TFCE technique. This work was supported by the Brazilian National Council for Scientific and Technological Development (CNPq), the Brazilian National Council for the Improvement of Higher Education (CAPES), the Swiss National Science Foundation (BSSG10_155915, 100014_178841, 32003B_166566, and PP00P1_170506/1), the Foundation for Research in Science and the Humanities at the University of Zurich (STWF-17-012), the Baugarten Stiftung, and the Swiss Government.

## Data and Code Availability Statement

All obtained results and scripts used for the data analysis are available on the public GitHub repository: https://github.com/gustavopamplona/Followup_NF_attention.

## Ethics Approval Statement

This study was approved by the local ethics committee of the Canton of Zurich in Switzerland. All participants read and signed the informed consent in accordance with the Declaration of Helsinki (2013) before taking part in the study.

