## Supplemental Figure 3 for "Long-term effects of network-based fMRI neurofeedback training for sustained attention"

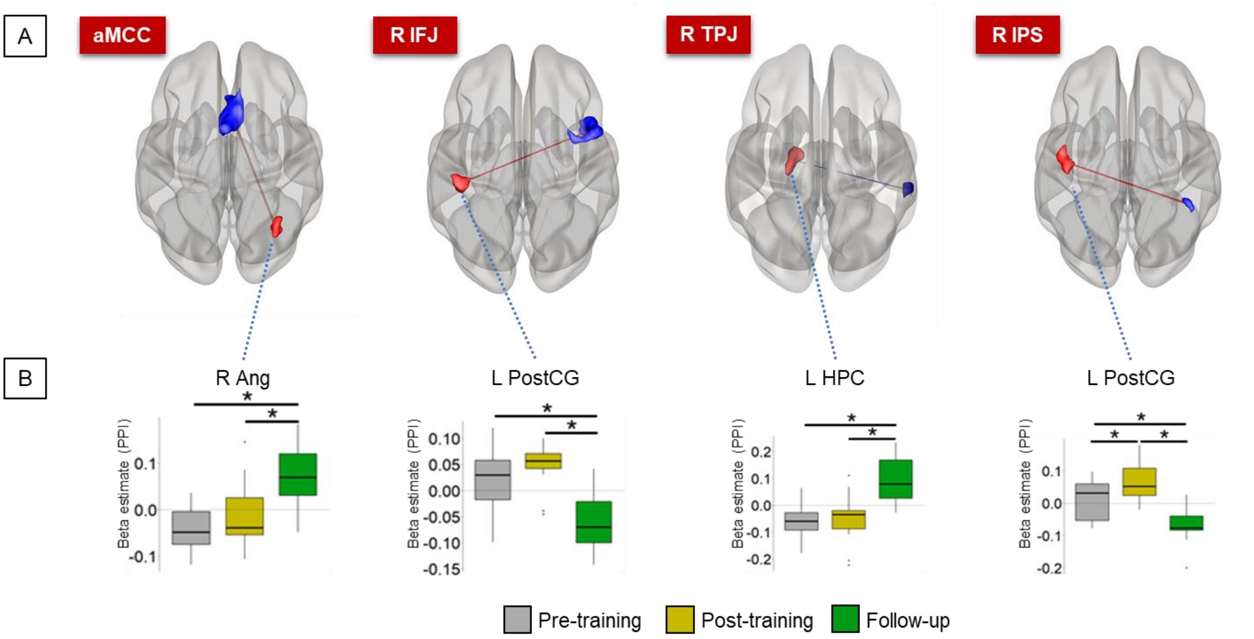


***Figure S1***. *(A) The regFC analyses showed that some clusters presented differences of FC with individual SAN regions across sessions, considering transfer runs. Blue and red clusters represent SAN regions and seed-to-voxel significant FC clusters, respectively, projected onto glass brains in superior view. (B) Boxplots represent the individual betas estimated for the PPI regressor of the SAN ROIs for each session; gray, yellow, and green represent pre-training, post-training, and follow-up sessions, respectively. The dashed black lines in the boxplots represent the zero level. Asterisks represent significant differences corrected for multiple comparisons by the Tukey method ($p < 0.05$). Ang = angular gyrus, PostCG = postcentral gyrus, HPC = hippocampus, L/R = left/right.*
