## Supplemental Figure 2 for "Long-term effects of network-based fMRI neurofeedback training for sustained attention"

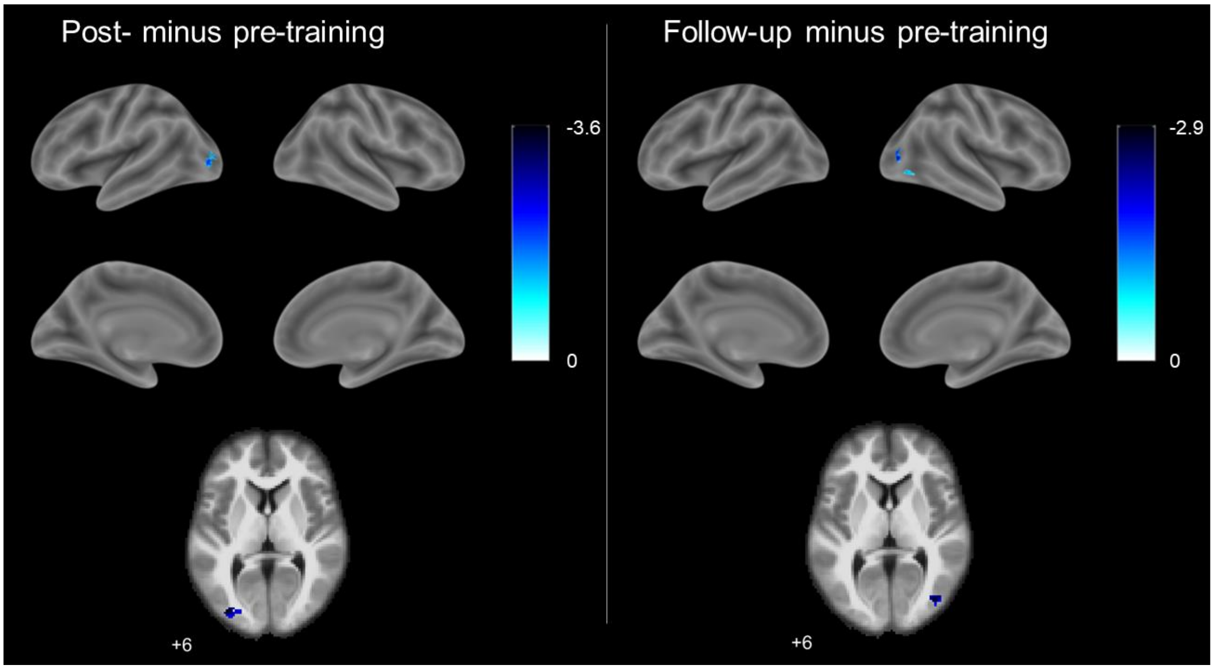


***Figure S2.*** *Whole-brain analyses show that the middle and inferior occipital cortex (part of the dorsal attention network) were less activated in post-training and follow-up sessions, compared to the pre-training session. Left and right columns show contrasts post- minus pre-training sessions and follow-up minus pre-training sessions, respectively. Cold colors represent significant negative t-values, respectively, overlapped onto surface-rendered (top) and axial slices (bottom) of a brain template.*
