## Supplemental Figure 1 for "Long-term effects of network-based fMRI neurofeedback training for sustained attention"

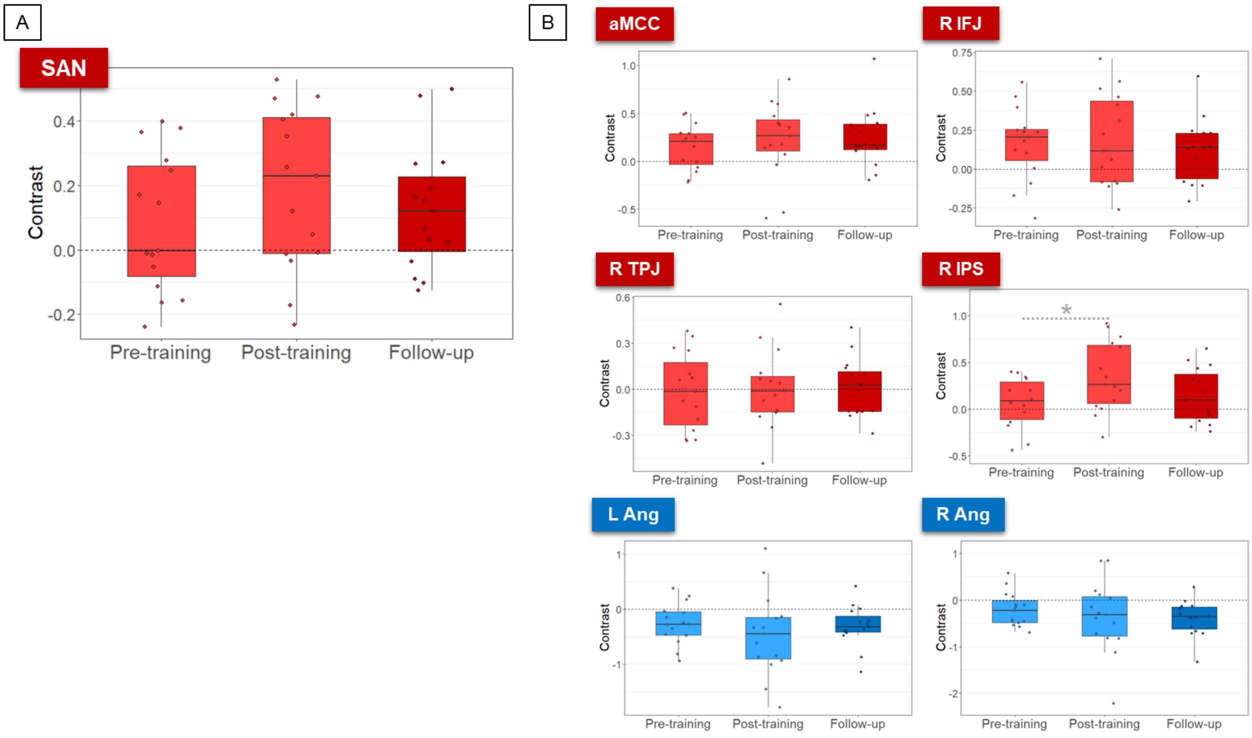


***Figure S1.*** *There were no significant differences in the activation of (A) SAN as an entity and (B) some SAN/DMN ROIs analyzed (aMCC = anterior midcingulate gyrus, R IFJ = right inferior frontal gyrus, R TPJ = right temporoparietal gyrus, R IPS = right intraparietal sulcus, L Ang = left angular gyrus, R Ang = right angular gyrus) across sessions of transfer runs. Light and dark colors represent pre-/post-training and follow-up sessions, respectively.*
