## Supplemental Table 2 for "Long-term effects of network-based fMRI neurofeedback training for sustained attention"

***Table S2.*** *Strategies for regulation and baseline blocks, as well as the concentration scores, reported by each participant during pre-training, post-training, and follow-up transfer runs.*

| Participant ID | Regulation/baseline strategies | | | Self-reported concentration scores (pre-training, post-training, follow-up noFB sessions) |
|---|---|---|---|---|
| | Pre-training noFB session | Post-training noFB session | Follow-up noFB session | |
| 1 | Attentional focus on the geometry of the up-arrow/Wandering thoughts, thinking about music | Attentional focus on the geometry of the up-arrow/Relaxation based on breathing | Attentional focus on the geometry of the up-arrow/Voluntary eye movements | 7, 9, 7 |
| 2 | Mental description the geometry of the up-arrow/Abandoned regulation strategy | Mental description the geometry of the up-arrow/Recalled past memories | Mental description the geometry of the up-arrow/Recalled past memories | 10, 7, 6.5 |
| 3 | Attentional focus on the geometry of the up-arrow/Thoughts about pleasant memories | Recalled past memories/Gazed deeply into the screen | Recalled past memories, future projections/Thought about nothing in particular | 6.5, 6, 6 |
| 4 | Tried to memorize the arrow, thought about the color of the arrow/Just looked at the squared without thinking about anything in particular | Counted the edges of the arrow/ Thoughts about the future, specifically about the next holidays | Focus on the geometry of the arrow/ Thoughts about the future | 6, 3, 8 |
| 5 | Attentional focus on the geometry of the up-arrow/Mental relaxation | Attentional focus on the geometry of the up-arrow/Thinking of sport events | Attentional focus on the geometry of the up-arrow/Thinking of sport events | 8, 7, 7 |
| 6 | Attentional focus on the geometry of the up-arrow/Focus on the light | Attentional focus on the geometry of the up-arrow/Random thoughts about his personal life | Attentional focus on the geometry of the up-arrow/Random thoughts about his personal life | 7, 7, 6.5 |
| 7 | Attentional focus on the geometry of the up-arrow/Not thinking about something specific | Mental math/Not thinking about something specific | Mental math/Not thinking about something specific | 7, 3, 4.5 |
| 8 | Mental simulation of schoolwork/Thought about nature | Mental simulation of schoolwork/Mental imagery of swimming | Mental simulation of schoolwork/Mental imagery of swimming | 8, 6, 6.5 |
| 9 | Focus on the geometry of the arrow and verbal associations/Mental focus on breathing | Mental translation of the arrow attributes into other languages/Mental focus on breathing | Mental translation of the arrow attributes into other languages/Mental focus on breathing | 7, 7, 5 |
| 10 | Attentional focus on the geometry of the up-arrow/blurred vision, singing songs in mind | Attentional focus on the geometry of the up-arrow/Thinking about cats and his girlfriend | Attentional focus on the geometry of the up-arrow/Mind-wandering | 7, 7, 6 |
| 11 | Imagery of a verbal fight/thought about falling asleep | Mental math/Mental focus on breathing | Mental math/Thought about nothing in particular | 5, 8, 8 |
| 12 | Focus on the geometry of the arrow and tried to keep a clear vision/Mental relaxation | Attentional focus on the geometry of the up-arrow/Thinking of a memory task | Imagery of running towards the arrow, imagery of a dangerous situation/Mental relaxation and auditory focus on the MRI sound | 7, 9, 6.5 |
| 13 | Attentional focus on the geometry of the up-arrow/Random thought and memories | Attentional focus on the geometry of the up-arrow; counting the edges of the up-arrow/Not thinking about something specific, trying to not focus vision on the square | Attentional focus on the geometry of the up-arrow; counting the edges of the up-arrow/Not thinking about something specific | 5, 7, 6 |
| 14 | Imagined the arrow as being a devil/Random thoughts | Mental imagery of giving a speech in front of a large audience/Recalling memories from the past; mental imagery of different pictures | Mental imagery of giving a speech in front of a large audience/Random thoughts | -, 10, 10 |
| 15 | Strategies not recorded (technical problem with the scanner communication system) | | | |
