## Supplemental Table 1 for "Long-term effects of network-based fMRI neurofeedback training for sustained attention"

***Table S1.*** *Selected SAN and DMN regions for the neurofeedback training.*

| Brain region | Number of voxels | MNI coordinates (mm) | | |
|---|---|---|---|---|
| | | x | y | z |
| **Anterior midcingulate cortex (aMCC)** | 682 | 2.0 | 14.1 | 47.0 |
| **Right inferior frontal junction (rIFJ)** | 345 | 46.8 | 7.1 | 34.5 |
| **Right temporoparietal junction (rTPJ)** | 85 | 60.8 | -33.9 | 13.7 |
| **Right intraparietal sulcus (rIPS)** | 29 | 43.9 | -45.6 | 46.6 |
| **Posterior cingulate cortex (PCC)** | 123 | 0 | -54 | 32 |
| **Medial prefrontal cortex (mPFC)** | 123 | 0 | 52 | 38 |
| **Left angular gyrus (lAng)** | 123 | -48 | -64 | 34 |
| **Right angular gyrus (rAng)** | 123 | 52 | -62 | 34 |

*Note: The SAN regions were the same for all subjects and the clusters were obtained from* (Langner and Eickhoff, 2013)*. The DMN regions were individually defined using resting-state scans and the Personode toolbox* (Pamplona et al., 2020b)*. MNI coordinates for DMN regions are shown as the median across participants on each axis.*
